# Osteocyte State Transitions Modulate Bone Remodeling During Early Skeletal Aging

**DOI:** 10.64898/2026.08.07.743422

**Authors:** Ryosuke Denda, Anhao Liu, Mikihito Hayashi, Cheng Wang, Haruhiko Akiyama, Hiroshi Takayanagi, Mitsuru Saito, Tomoki Nakashima

**Affiliations:** Department of Cell Signaling, Graduate School of Medical and Dental Sciences, Institute of Science Tokyo, Tokyo, Japan, 113-8549; Department of Orthopaedic Surgery, The Jikei University School of Medicine, Tokyo, Japan,105-8461; Department of Biochemistry, School of Dentistry, Showa Medical University, Tokyo, Japan, 142-8555; Department of Orthopaedic Surgery, Gifu University School of Medicine, Gifu, Japan, 501-1194; Center for One Medicine Innovative Translational Research (COMIT), Institute for Advanced Study, Gifu University, Gifu, Japan, 501-1194; Department of Immunology, Graduate School of Medicine and Faculty of Medicine, The University of Tokyo, Tokyo, Japan, 113-0033; Faculty of Dentistry, Institute of Science Tokyo, Tokyo, Japan, 113-8549

## Abstract

Osteocytes are long-lived cells that play a central role in bone homeostasis, yet age-related changes in their functional states remain poorly understood, particularly because skeletal aging involves multiple processes beyond cellular senescence. We generated an osteocyte-specific *Mepe*^Cre^ mouse line and combined osteocyte ablation in young and middle-aged mice with skeletal phenotyping, single-cell transcriptomics, and senolytic treatment. *Mepe*^Cre^-driven recombination was largely confined to osteocytes, with minimal off-target activity. Osteocyte ablation increased bone mass at both ages, indicating that osteocytes constrain bone accrual as part of their role in skeletal homeostasis. However, the accompanying remodeling changes differed with age: enhanced osteoblast activity predominated in young mice, whereas reduced osteoclast-mediated bone resorption predominated in middle-aged mice. Single-cell transcriptomics revealed distinct osteocyte subpopulations whose relative abundance shifted with age, from a predominantly matrix-enriched state in young mice to an expanded aging-transitional state in middle-aged mice. Although this state showed partial enrichment of senescence-associated transcriptional signatures, senolytic treatment failed to recapitulate the increase in bone mass induced by osteocyte ablation. Osteocyte therefore regulate bone mass through age-dependent mechanisms that coincide with shifts in osteocyte-state composition. These changes emerge by middle age and may contribute to early remodeling imbalance before overt cellular senescence during skeletal aging.

**Graphical Abstract:** Graphical summary of the findings of this study. AA, amino acids; NA, nucleic acid; UA, uric acid; TCA, tricarboxylic acid.

## Introduction

Osteocytes are long-lived cells embedded within the bone matrix and serve as central regulators of bone homeostasis(1, 2). Several studies, including genetic ablation models such as *Dmp1* promoter-driven diphtheria toxin receptor (DTR) systems, have demonstrated that osteocytes regulate osteoclast and osteoblast activity through the production of soluble and membrane-bound factors, including osteoprotegerin (OPG)(3), receptor activator of nuclear factor-κB ligand (RANKL)(4, 5), and semaphorin 3A(6)— and even exert a systemic influence via sclerostin (encoded by the *Sost* gene)(7, 8) and fibroblast growth factor 23 (FGF23)(9). These insights have largely been derived from approaches that treat osteocytes as a relatively uniform cell population. However, recent advances in single-cell RNA sequencing (scRNA-seq) have revealed previously unrecognized heterogeneity among osteocytes(10), raising the possibility that distinct osteocyte states contribute differentially to bone homeostasis, particularly during aging.

Osteocytes have been implicated in the process of skeletal aging due to their long lifespans(11–13), and recent studies have shown that osteocyte senescence contributes to bone loss(14). While systemic clearance of senescent cells can alleviate age-related bone loss, local elimination of senescent osteocytes appears less effective in restoring bone mass(15), suggesting that factors beyond osteocyte senescence—or contributions from other cell types—may underlie skeletal aging. Aging is a continuous process, and the changes that occur prior to the establishment of overt senescence remain poorly defined(16, 17). In particular, it is unclear whether osteocytes undergo functional or phenotypic transitions during these intermediate stages, and how such changes affect bone remodeling. The limitation of existing genetic tools have hampered efforts to address this question. The widely used *Dmp1*–Cre-based models, although informative in earlier osteocyte studies, exhibit recombination activity in non-skeletal tissues and osteoblasts, raising concerns about targeting specificity(18–20). Alternative Cre drivers, such as *Sost*–Cre and *Sost*–Cre^ERT2^, have been developed but remain insufficient for the exclusive targeting of osteocytes(21, 22).

Here, we developed an improved osteocyte-specific Cre driver, *Mepe*^Cre^, leveraging the restricted expression profile of *Mepe* in the osteocyte lineage. Using this model, we found that osteocyte ablation increased bone mass in both young and middle-aged mice, indicating that osteocytes restrict bone mass under physiological conditions, contrary to the prevailing view. To explore the mechanisms underlying these age-dependent differences, we performed scRNA-seq on isolated osteocytes, which revealed heterogeneous osteocyte subpopulations with dynamic shifts. In particular, a matrix-enriched subpopulation associated with extracellular matrix production and OPG expression predominated in young mice, whereas an aging-transitional subpopulation associated with RANKL and inflammatory cytokine expression accumulated with aging. These state shifts provide a potential mechanistic basis for the age-dependent effects of osteocytes on bone mass. Consistently, the limited efficacy of senolysis in middle-aged mice, together with low p16 expression in the aging-transitional subpopulation, suggests that osteocyte state transitions may precede overt senescence.

## Results

### Osteocyte ablation using *Mepe*^Cre^ induces an increase in bone mass in young mice

To enable precise genetic targeting of osteocytes, we generated a knock-in mouse line in which an IRES–Cre cassette was inserted into the endogenous *Mepe* locus (*Mepe*^Cre^), based on its restricted expression pattern(23),(24) (Figs. 1A, S1A, S1B). Reporter analyses using this mouse line confirmed that *Mepe*^Cre^ activity was largely restricted to osteocytes in bone, with minimal activity in osteoblasts, chondrocytes or non-skeletal tissues, indicating improved osteocyte specificity compared to previously used Cre drivers (Figs. 1B–D, Fig. S1C, D).

**Figure 1.**
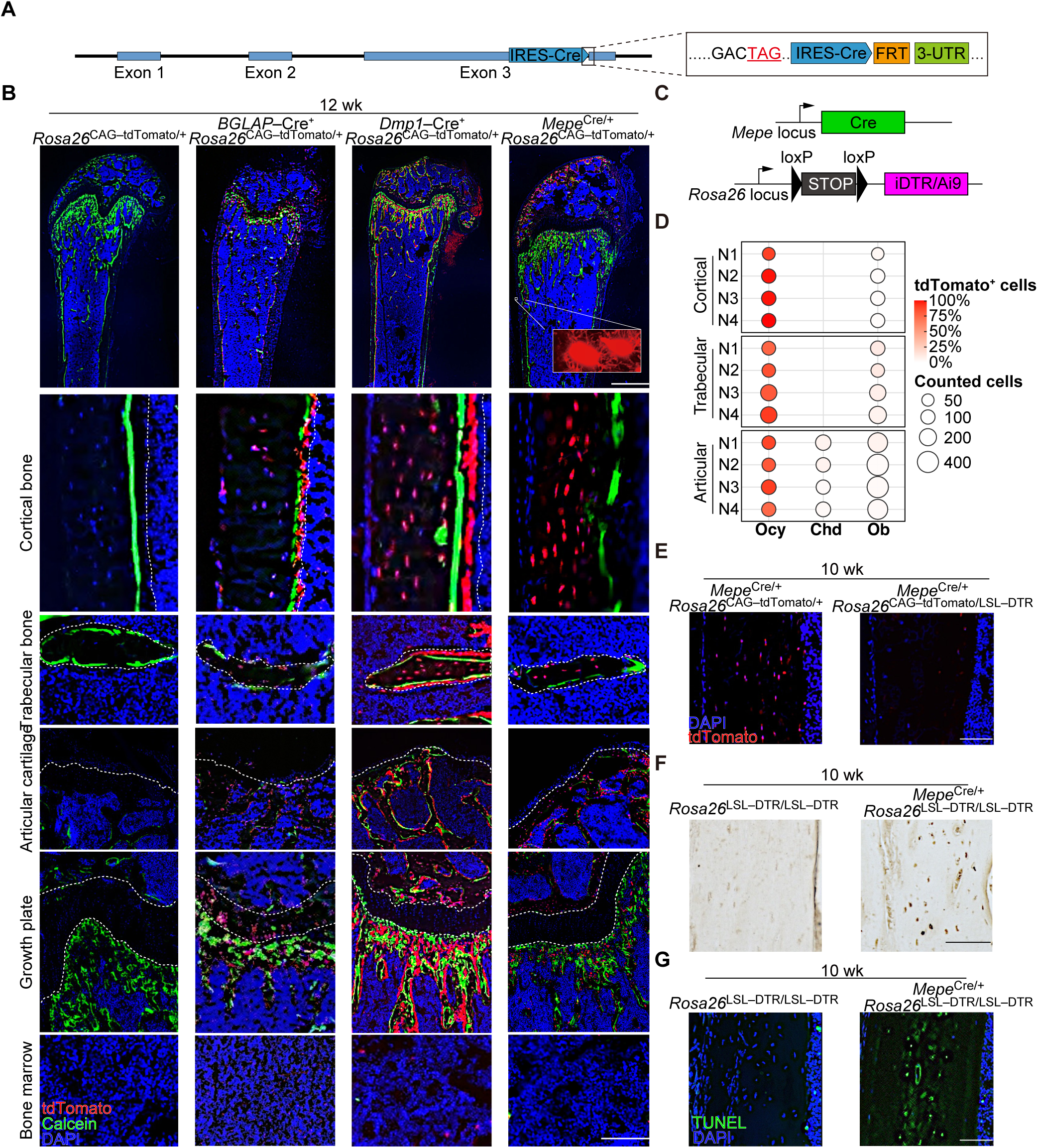
Improved specificity of osteocyte targeting and ablation using *Mepe*^Cre^. **(A)** Design of the *Mepe*^Cre^ allele. (**B)** Schematic representation of the allele carried by *Mepe*^Cre/+^ *Rosa26*^LSL–DTR/LSL–DTR^ or *Mepe*Cre/+ *Rosa26*^CAG–tdTomato^/+ mice. **(C)** Representative images of frozen sections of distal femur from male *Mepe*^Cre/+^ *Rosa26*^CAG–tdTomato/+^ and *Mepe*^Cre/+^ *Rosa26*^CAG–tdTomato/LSL–DTR^ mice. Red, tdTomato; green, calcein; blue, DAPI. Dotted lines indicate tissue boundaries: bone vs. bone marrow (Cortical and trabecular bone panel), bone vs. growth plate (Growth plate panel), and bone vs. articular cartilage (Articular cartilage panel). All specimens were obtained from 10-week-old male mice. Scale bars, 1 mm (low magnification) and 100 μm (high magnification). (**D)** Regional quantification of tdTomato labeling in femurs from *Mepe*^Cre/+^ *Rosa26*^CAG–tdTomato/+ mice^. Bubble color indicates the percentage of tdTomato-positive cells, and bubble size represents the number of cells counted as the denominator for each parameter. Ocy (osteocytes), tdTomato-positive osteocytes among all osteocytes; Chd (chondrocytes), tdTomato-positive chondrocytes among all chondrocytes; Ob (osteoblasts), tdTomato-positive cells that completely overlap with calcein labeling among all tdTomato-positive cells. Each row represents one mouse. (**E)** Representative images of frozen sections of distal femoral cortical bone from male *Mepe*^Cre/+^ *Rosa26*^CAG–tdTomato/+^ and *Mepe*^Cre/+^ *Rosa26*^CAG–tdTomato/LSL–DTR^ mice at 3 weeks after DT injection. Red, tdTomato. Scale bars, 100 μm. (**F)** Representative images of immunohistochemical staining of diphtheria toxin receptor (DTR) in tibiae from *Rosa26*LSL–DTR/LSL–DTR and *Mepe*^Cre/+^ *Rosa26*^LSL–DTR/LSL–DTR^ mice. Scale bar, 100 μm. (**G)** Representative images of TUNEL staining of femoral cortical bone from 10-week-old male *Rosa26*^LSL–DTR/LSL–DTR^ and *Mepe*^Cre/+^ *Rosa26*^LSL–DTR/LSL–DTR^ mice at 3 weeks after DT injection. Green, TUNEL; blue, DAPI. Scale bar, 100 μm.

We next induced osteocyte-specific ablation by crossing *Mepe*^Cre^ mice with inducible diphtheria toxin receptor (iDTR) mice (Figs. 1B). Efficient ablation was confirmed by the marked reduction of reporter-positive osteocytes and the appearance of empty lacunae in cortical bone (Figs. 1E–G). Because a pilot study revealed prominent empty osteocyte lacunae 3 weeks after the final diphtheria toxin (DT) injection, this time point was used for all subsequent analyses (Figs. 2A, B). Strikingly, osteocyte ablation in young mice resulted in a significant increase in both trabecular and cortical bone mass, accompanied by a significant decrease in serum sclerostin and FGF23 (Figs. 2C–E).

**Figure 2.**
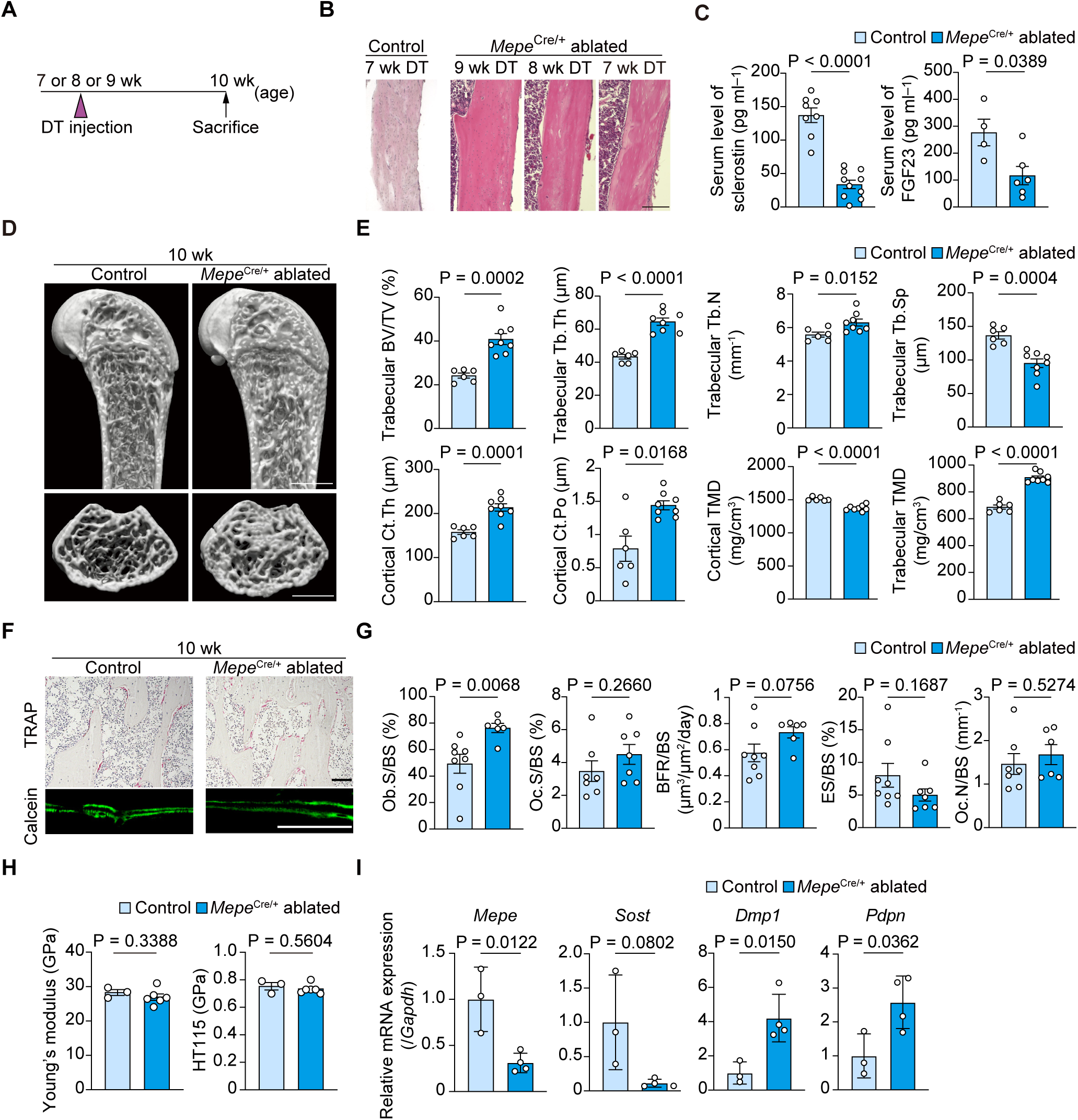
Osteocyte ablation using *Mepe*^Cre^ increases bone mass in young mice. **(A)** Experimental schedule of short-term ablation of *Mepe*-expressing osteocytes. Diphtheria toxin (DT) was administered 1, 2, or 3 weeks prior to analysis. (**B)** Representative histological images of proximal tibiae from 10-week-old *Rosa26*^LSL–DTR/LSL–DTR^ and *Mepe*^Cre/+^ *Rosa26*^LSL–DTR/LSL–DTR^ mice following DT injection at indicated time points. Scale bars, 200 μm. (**C, D)** μCT analysis of distal femurs from 10-week-old male *Rosa26*^LSL–DTR/LSL–DTR^ and *Mepe*^Cre/+^ *Rosa26*^LSL–DTR/LSL–DTR^ mice at 3 weeks after DT injection (n = 6–8 mice per group; scale bars, 1 mm. (**E)** Serum levels of sclerostin and FGF23 in 10-week-old male *Rosa26*^LSL–DTR/LSL–DTR^ and *Mepe*^Cre/+^ *Rosa26*^LSL–DTR/LSL–DTR^ mice (n = 4–10 mice per group) 3 weeks after DT injection. (**F)** Bone histomorphometric analysis of proximal tibiae from 10-week-old male *Rosa26*^LSL–DTR/LSL–DTR^ and *Mepe*^Cre/+^ *Rosa26*^LSL–DTR/LSL–DTR^ mice (n = 6–8 mice per group) 3 weeks after DT injection; Scale bars, 100 μm. (**G)** Relative *Mepe*, *Sost*, *Dmp1*, and *Pdpn* mRNA expression in bone of 10-week-old male *^Rosa26^*^LSL–DTR/LSL–DTR^ and *Mepe*^Cre/+^ *Rosa26*^LSL–DTR/LSL–DTR^ mice (n = 3–4 mice per group) 3 weeks after DT injection. (**H)** Biomechanical properties of femurs from 10-week-old male *Rosa26*^LSL–DTR/LSL–DTR^ and *Mepe*^Cre/+^ *Rosa26*^LSL–DTR/LSL–DTR^ mice (n = 3–6 mice per group) 3 weeks after DT injection, evaluated by nanoindentation analysis. All data are presented as mean ± SEM. Statistical significance was determined by Welch’s *t*-test after excluding outliers via Grubbs’ test.

Histomorphometric analysis further showed that this increase was accompanied by elevated osteoblast activity along with sclerostin downregulation, whereas changes in osteoclast parameters were relatively modest (Fig. 2F). To exclude the possibility that this increase in bone mass was due to unintended ablation of osteoblast-lineage cells, we administered DT to *BGLAP*-Cre; iDTR mice. In contrast to *Mepe*^Cre^-mediated osteocyte ablation, *BGLAP*-Cre-mediated ablation resulted in reduced bone mass (Figs. S2A, B), indicating that the bone gain phenotype observed in *Mepe*^Cre^ mice is not attributable to osteoblast loss. The gene expression pattern of the osteocyte lineage further supported this finding. Expression of mature osteocyte markers, including *Mepe* and *Sost*, decreased after ablation, whereas expression of early osteocyte markers, such as *Dmp1* and *Pdpn*, increased, indicating that new bone matrix formation by osteoblasts predominated during this process (Fig. 2G). However, the mechanical properties of bone shows no significant improvement, likely due to a concurrent increase in porosity (Fig. 2H).

### Age-dependent differences in bone remodeling following osteocyte ablation

To determine how the skeletal effects of osteocyte ablation change over time, we examined longitudinal changes in young mice following a single DT injection. µCT analysis showed that trabecular bone mass remained elevated for several weeks after ablation but gradually declined thereafter (Fig. 3). In cortical bone, the increase in thickness was more transient, whereas bone mineral density remained elevated for a longer period. These findings indicate that the effects of osteocyte ablation are dynamic and change with age.

**Figure 3.**
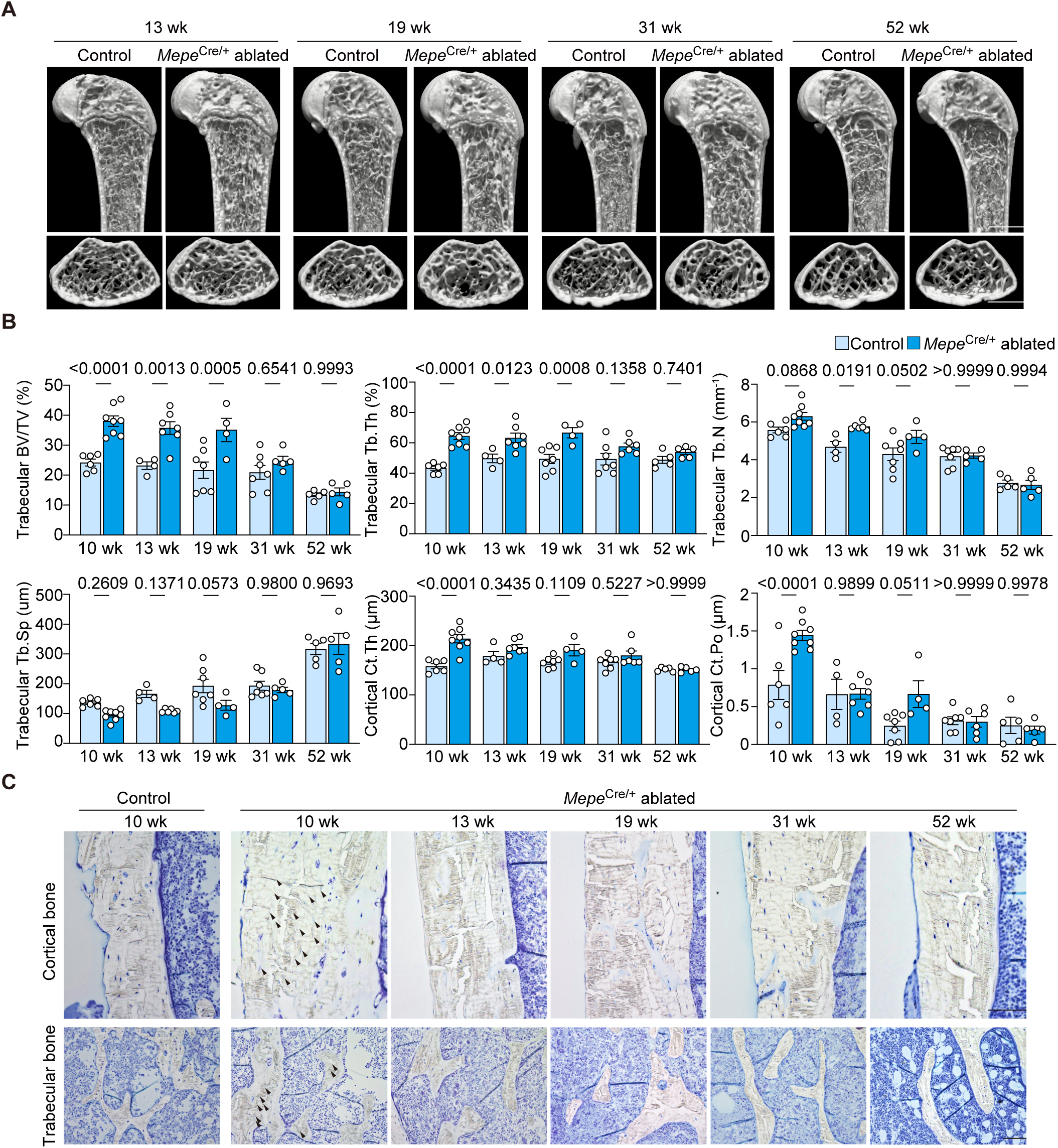
*Mepe*^Cre^-driven osteocyte ablation induces a sustained activation of bone formation, which wanes with the aging process. **(A–C) A:** Representative μCT images; **B**: μCT analysis; **C**: representative toluidine blue staining images (of distal femurs from 10-, 13-, 19-, 31- and 52-week-old male *Rosa26*^LSL–DTR/LSL–DTR^ and *Mepe*^Cre/+^ *Rosa26*^LSL–DTR/LSL–DTR^ mice (n = 4–8 mice per group) following DT injection at 7 weeks of age. **A**: Scale bars, 1 mm; **C**: 100 μm. Data in **B** are presented as mean ± SEM. Statistical significance was determined by two-way ANOVA with Sidak’s test after excluding outliers via Grubbs’ test.

We next asked whether osteocyte ablation exerts similar effects at later stages of adulthood. In 52-week-old mice, osteocyte ablation also resulted in a significant increase in both trabecular and cortical bone mass, accompanied by sclerostin and FGF23 downregulation (Figs. 4A–D), indicating that the paradoxical bone gain is maintained with age during adulthood. However, histomorphometric analysis revealed that the underlying remodeling dynamics differed substantially between age groups. In young mice, bone gain was primarily driven by enhanced osteoblast activity, with osteoclast parameters largely unchanged (Figs. 2F, G). In contrast, in middle-aged mice, osteocyte ablation led to a marked reduction in osteoclast number and activity, with comparatively modest changes in osteoblast-related parameters (Fig. 4E). Meanwhile, the increased expression of early osteocyte markers was no longer observed, indicating that the increase in bone mass in middle-aged mice was independent of enhanced osteogenesis (Fig. S3A).

**Figure 4.**
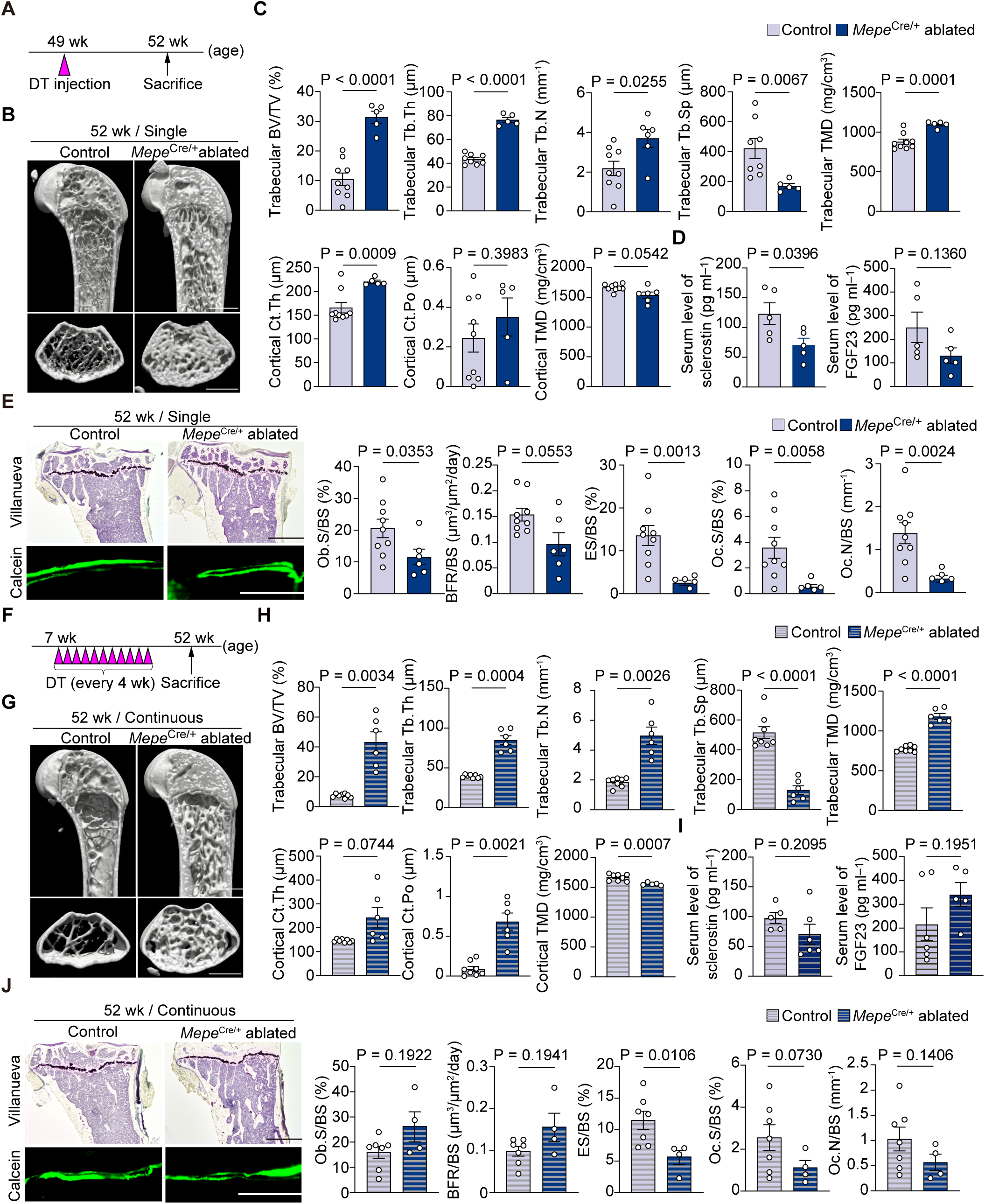
*Mepe*^Cre^-mediated osteocyte ablation increases bone mass in middle-aged mice by suppressing osteoclast activity. **(A, F)** Experimental schedule for the ablation of *Mep*e-expressing osteocytes in 1-year-old mice. DT was administered 3 weeks before the analysis or monthly. **(B, C**, **G, H)** μCT analysis of distal femurs from 1-year-old male *Rosa26*^LSL–DTR/LSL–DTR^ and *Mepe*^Cre/+^ *Rosa26*^LSL–DTR/LSL–DTR^ mice that received a single DT injection (**B** and **C**; n = 5–9 mice per group) or continuous DT injections (**G** and **H**; n = 6–8 mice per group); Scale bars, 1mm. (**D, I)** Serum sclerostin and FGF23 concentrations in 52-week-old male *Rosa26*^LSL–DTR/LSL–DTR^ and *Mepe*^Cre/+^ *Rosa26*^LSL–DTR/LSL–DTR^ mice (n = 4–10 mice per group) 3 weeks after a single DT injection or after the final injection of the continuous regimen. (**E**, **J)** Bone histomorphometric analysis of distal femurs from 1-year-old male *Rosa26*^LSL–DTR/LSL–DTR^ and *Mepe*^Cre/+^ *Rosa26*^LSL–DTR/LSL–DTR^ mice that received a single DT injection (**E**; n = 5–9 mice per group) or continuous DT injections (**J**; n = 4–7 mice per group); Scale bars, 500 μm (top) and 100 μm (bottom). All data are presented as mean ± SEM. Statistical significance was determined by Welch’s *t*-test after excluding outliers via Grubbs’ test.

To determine whether these age-dependent differences reflect the cumulative consequences of aging or an intrinsic change in osteocyte function, we performed sustained osteocyte ablation by repeatedly administering DT from a young age (Fig. 4F).

Under these conditions, the high bone mass phenotype persisted, remaining driven by increased osteoblast activity, as in young mice, despite the absence of significant changes in serum sclerostin and FGF23 levels (Figs. 4G–J, S3B). Together, these findings indicate that osteocyte ablation increases bone mass across ages but through distinct remodeling mechanisms, suggesting that the functional properties of osteocytes—and thus the consequences of their removal—shift with age.

### Distinct osteocyte states are dynamically restructured with age

To define osteocyte states across adulthood, we performed scRNA-seq on highly purified *Mepe*-expressing osteocytes isolated from 12- and 52-week-old reporter mice (Figs. 5A, B). Following quality control, removal of hematopoietic-lineage cells, and osteocyte gene set-based subclustering(8), UMAP analysis resolved the osteocyte-enriched populations into 18 transcriptionally distinct clusters, revealing substantial heterogeneity across different life stages (Figs. 5C–E). Stress-related gene sets (*Fos*, *Jun*, *Junb*, *Jund*, *Atf3*, *Egr1*, *Dusp1*, *Hspa1a*, *Hspa1b*, *Ier2*, *Ier3*, and *Zfp36*) and cell cycle scores were evaluated to ensure that the observed heterogeneity was not caused by the tissue digestion procedure (Fig. 5F).

**Figure 5.**
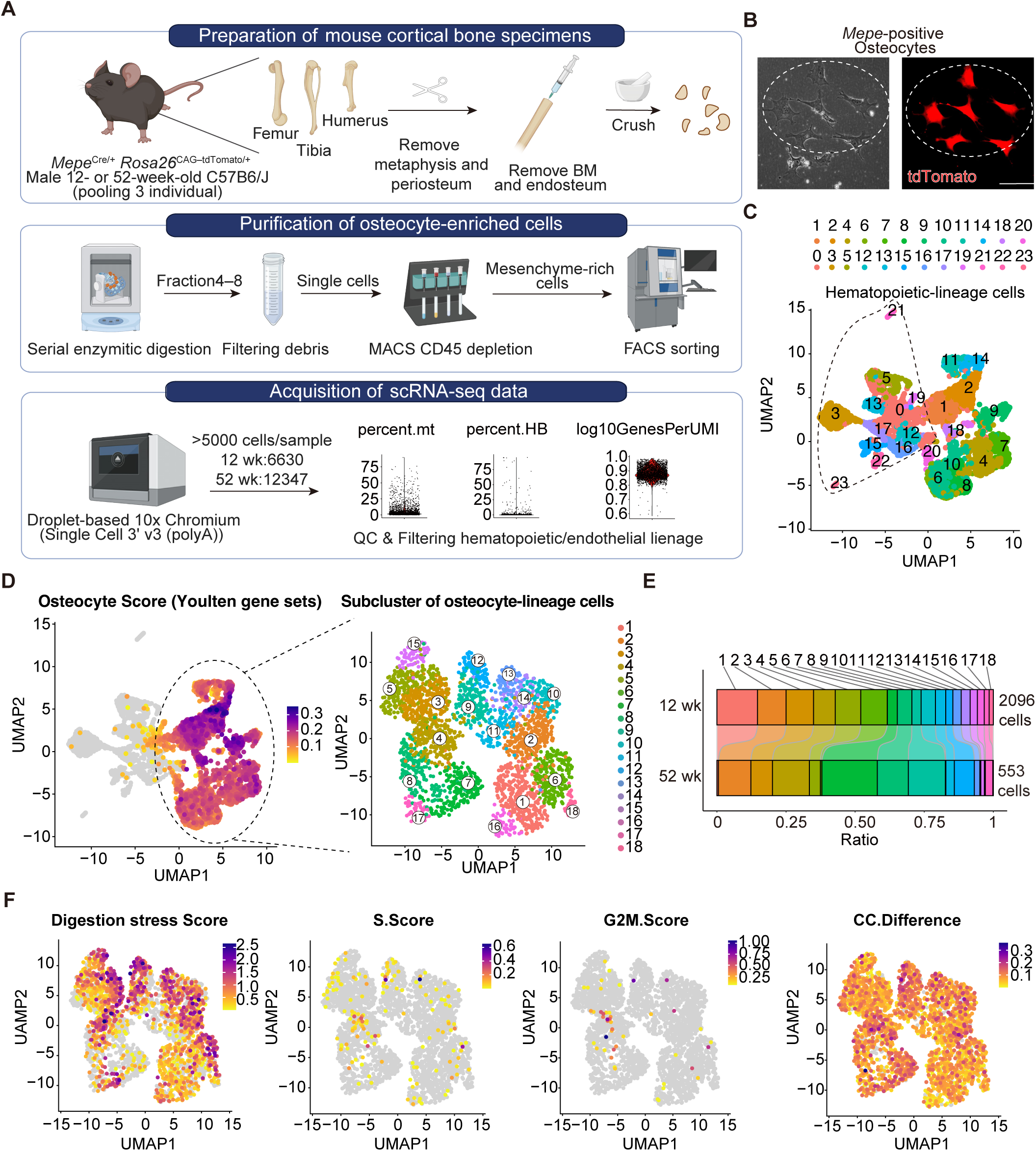
*Mepe*^Cre^-sorted osteocytes exhibit transcriptional heterogeneity. **(A)** Workflow for preparation of scRNA-seq samples from mouse long bones. **(B)** Representative fluorescence images of sorted osteocytes from *Mepe*^Cre/+^ *Rosa26*^CAG–tdTomato/+^ mice. Scale bar, 20 μm. **(C)** UMAP visualization of cell identities obtained by scRNA-seq. Clusters enclosed by the dotted line were identified as contaminating hematopoietic-lineage cells based on marker gene expression. Distinct cell populations are indicated by different colors. Each dot represents a single cell. **(D)** Visualization of gene set scoring based on the osteocyte gene set reported by Youlten *et al*. The UMAP of the subclusters after exclusion of hematopoietic-lineage cells is shown on the right side. These cells were defined as tdTomato-expressing osteocyte-lineage cells and were used for all subsequent analyses. **(E)** Distribution of clusters across the 12- and 52-week-old samples. **(F)** Visualization of gene set scoring based on stress-related and cell cycle-related gene sets. CC, cell cycle. CC.Difference = G2M.Score – S.score.

Given the graded expression pattern observed among the osteocyte clusters, we applied CytoTRACE(25) and RNA velocity(26) analyses to define the differentiation landscape of these populations (Fig. 6A). These analyses identified cluster 14 as the least differentiated population and supported a continuous differentiation trajectory (Figs. 6B, C). Pseudotime analysis using this cluster as a root revealed three major maturation trajectories, each terminating at a distinct endpoint (Fig. 6D). Based on trajectory architecture and distinct marker gene expression patterns, we defined four osteocyte subpopulations: canonical, matrix-enriched, aging-transitional, and immature (Figs. 6E, S4, S5). The aging-transitional subpopulation exhibited increased expression of remodeling-associated and inflammatory genes, whereas the matrix-enriched subpopulation was characterized by elevated expression of extracellular matrix–related genes, supporting their functional annotations. Notably, the relative abundance of these subpopulations changed markedly with age. The matrix-enriched subpopulation predominated in young mice, whereas the aging-transitional subpopulation expanded in middle-aged mice, indicating a substantial shift in subpopulation composition during aging (Fig. 6E).

**Figure 6.**
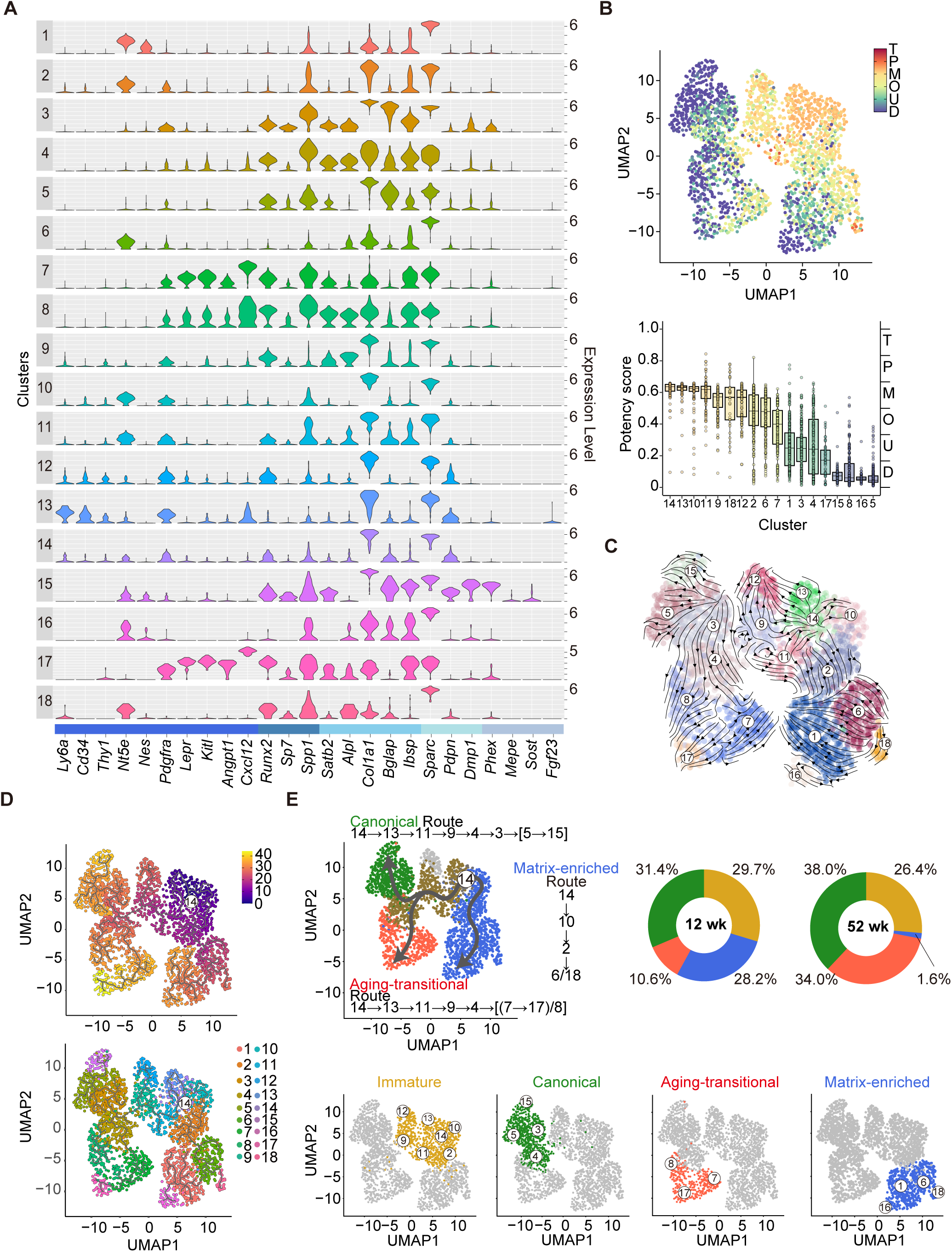
*Mepe*^Cre^-sorted osteocytes exhibit distinct states whose composition is dynamically restructured with age. **(A)** Violin plots showing the expression of mesenchymal-osteolineage-related markers among clusters. **(B)** Analysis of differentiation potential based on CytoTRACEv2. T, totipotent; P, pluripotent; M, multipotent; O, oligopotent; U, unipotent; D, differentiated. **(C)** RNA Velocity analysis based on scVelo. **(D)** Pseudotime trajectory analysis based on Monocle3. The root point was defined based on the results shown in **B** and **C**. **(E)** Maturation trajectories of distinct osteocyte subpopulations (upper left) and their corresponding Seurat cluster composition based on marker gene expression (lower). Quantification of the relative abundance of each subpopulation across ages (upper right).

These subpopulations were further supported by distinct metabolic signatures revealed by scFEA(10) analysis (Fig. S6A). The matrix-enriched subpopulation exhibited enhanced lipid biosynthesis, active glycolysis with lactate production, and increased protein glycosylation, whereas the aging-transitional subpopulation showed elevated amino acid catabolism, increased nucleotide turnover, and enhanced tricarboxylic acid (TCA) cycle activity (Figs. S6A, B). This metabolic divergence further supports the existence of functionally distinct osteocyte states. Together, these findings demonstrate that osteocytes comprise heterogeneous subpopulations organized along defined differentiation trajectories, and that their composition is dynamically restructured with age, providing a cellular basis for the life stage-dependent differences in bone remodeling.

### Osteocyte subpopulations exhibit divergent regulatory programs associated with bone remodeling

To determine the functional significance of these osteocyte subpopulations, we compared their gene expression profiles. The aging-transitional osteocytes showed increased expression of growth factor–related genes, including *Igfbp4* and *Igfbp5*, whereas the matrix-enriched osteocytes exhibited transcriptional features reminiscent of chondrocyte-like profiles, including the expression of extracellular matrix–related genes such as *Col2a1* and *Col9a2* (Fig. 7A). In contrast, canonical osteocytes retained higher expression of classical osteolineage markers such as *Runx2* and *Bglap*. Gene ontology analysis further indicated that, while these subpopulations shared core osteocytic features, they differed in membrane-associated signaling and extracellular interactions. The aging- transitional osteocytes showed increased responsiveness to growth factors and cell adhesion pathways, whereas the matrix-enriched osteocytes were enriched for glycan-associated extracellular processes (Fig. 7B).

**Figure 7.**
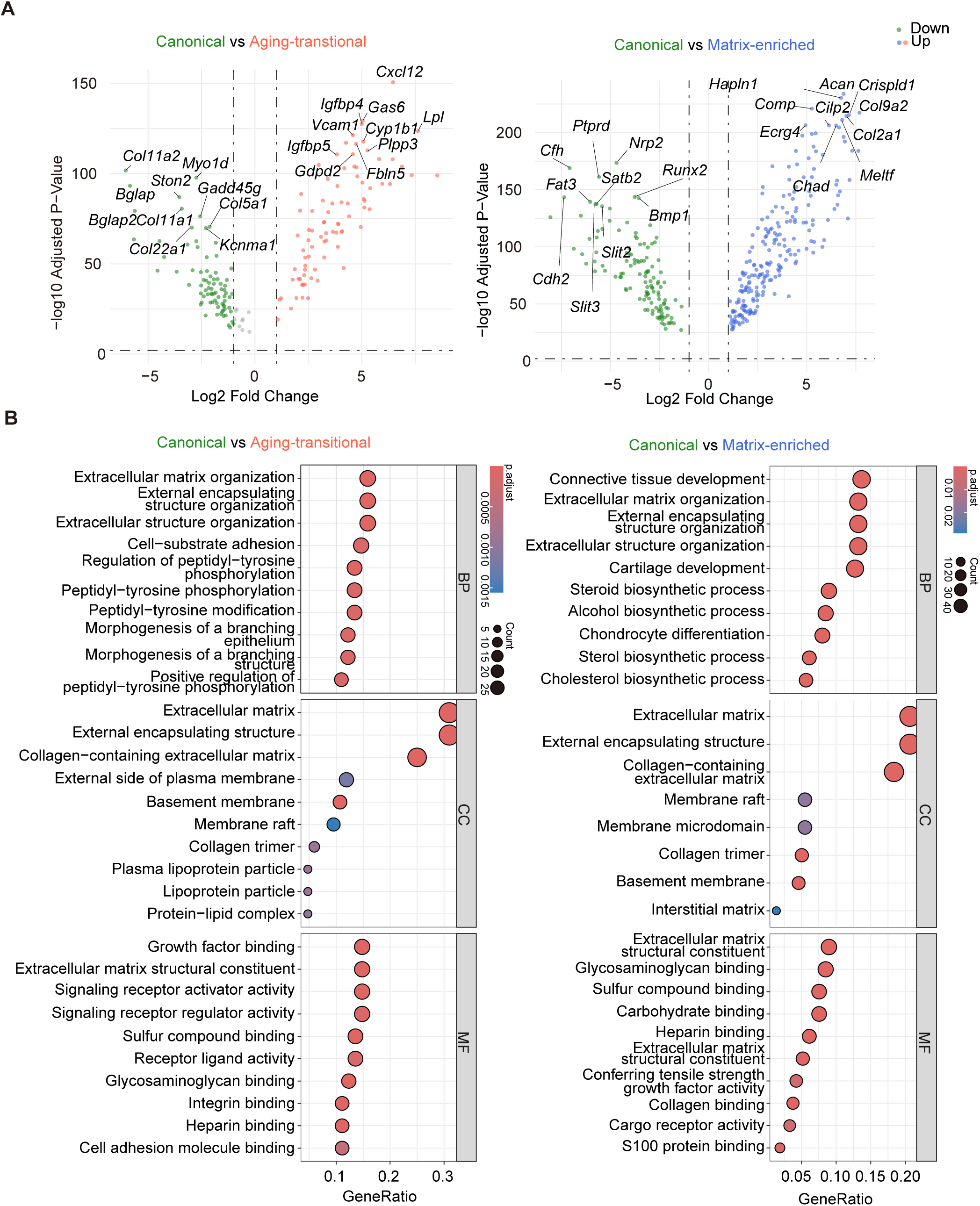
Osteocyte subpopulations exhibit divergent regulatory programs associated with bone remodeling. **(A)** Visualization of differentially expressed genes (DEGs) among the canonical and matrix-interacting/matrix-enriched subpopulations. Cutoffs: Log2 fold change ≥ 1; adjusted p-value < 0.01. **(B)** Pathway enrichment analysis of DEGs from (A), based on Gene Ontology (GO) database. BP, biological process; CC, cellular component; MF, molecular function.

Notably, these transcriptional differences were accompanied by distinct expression patterns of key regulators of bone remodeling, RANKL and OPG. The aging- transitional subpopulation showed increased expression of *Tnfsf11* (encoding RANKL), along with inflammatory cytokines, whereas the matrix-enriched subpopulation exhibited higher expression of *Tnfrsf11b* (encoding OPG; Figs. 8A, B). In contrast, the canonical subpopulation maintained relatively balanced expression of these factors. These findings indicate that osteocyte subpopulations are functionally distinct and are differentially linked to age-dependent remodeling outcomes. In particular, the expansion of the aging- transitional subpopulation is consistent with enhanced osteoclast-supporting potential through elevated RANKL expression and inflammatory signaling in middle-aged mice. In contrast, the matrix-enriched subpopulation is associated with extracellular matrix–related programs and a relative enrichment of OPG expression; however, these features alone do not fully account for the increased bone formation observed in young mice.

**Figure 8.**
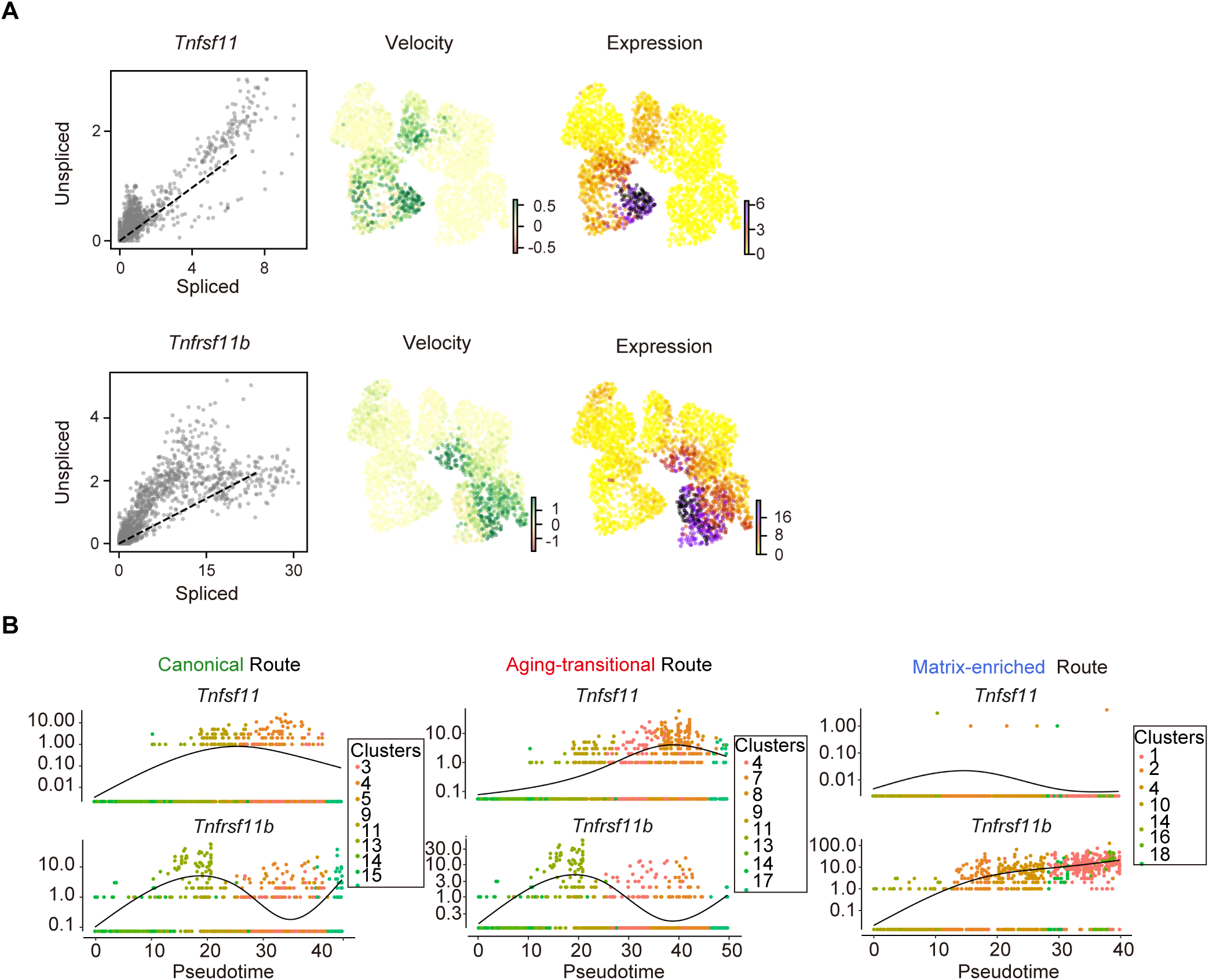
*Tnfsf11* and *Tnfrsf11b* expression changes across osteocyte subpopulations. **(A)** RNA velocity and expression levels of *Tnfsf11* and *Tnfrsf11b* analyzed by scVelo. **(B)** Pseudotime analysis of *Tnfsf11* and *Tnfrsf11b* across the distinct maturation trajectories using Monocle3.

### Osteocyte state transitions are early events that partially precede overt senescence

To determine whether the observed osteocyte state transitions are driven by cellular senescence, we quantitatively assessed senescence-associated signatures in our scRNA-seq dataset. Expression of canonical senescence markers, including *Cdkn2a* (p16) and *Cdkn1a* (p21), was detected in a limited subset of osteocytes and showed only partial enrichment within the aging-transitional subpopulation (Figs. 9A, B). Consistently, both scoring with the senescence-associated gene set (SenMayo(27)) and a comprehensive evaluation of senescence and aging using senePy and tAge revealed a modest but non-exclusive enrichment in this subpopulation, indicating that senescence-related transcriptional programs are present but insufficient to fully account for this state (Figs. 9C–E). Importantly, the elevation of senescence signatures did not fully overlap with the aging-transitional cluster, suggesting that this population represents a broader remodeling-associated state rather than a purely senescent cell population.

**Figure 9.**
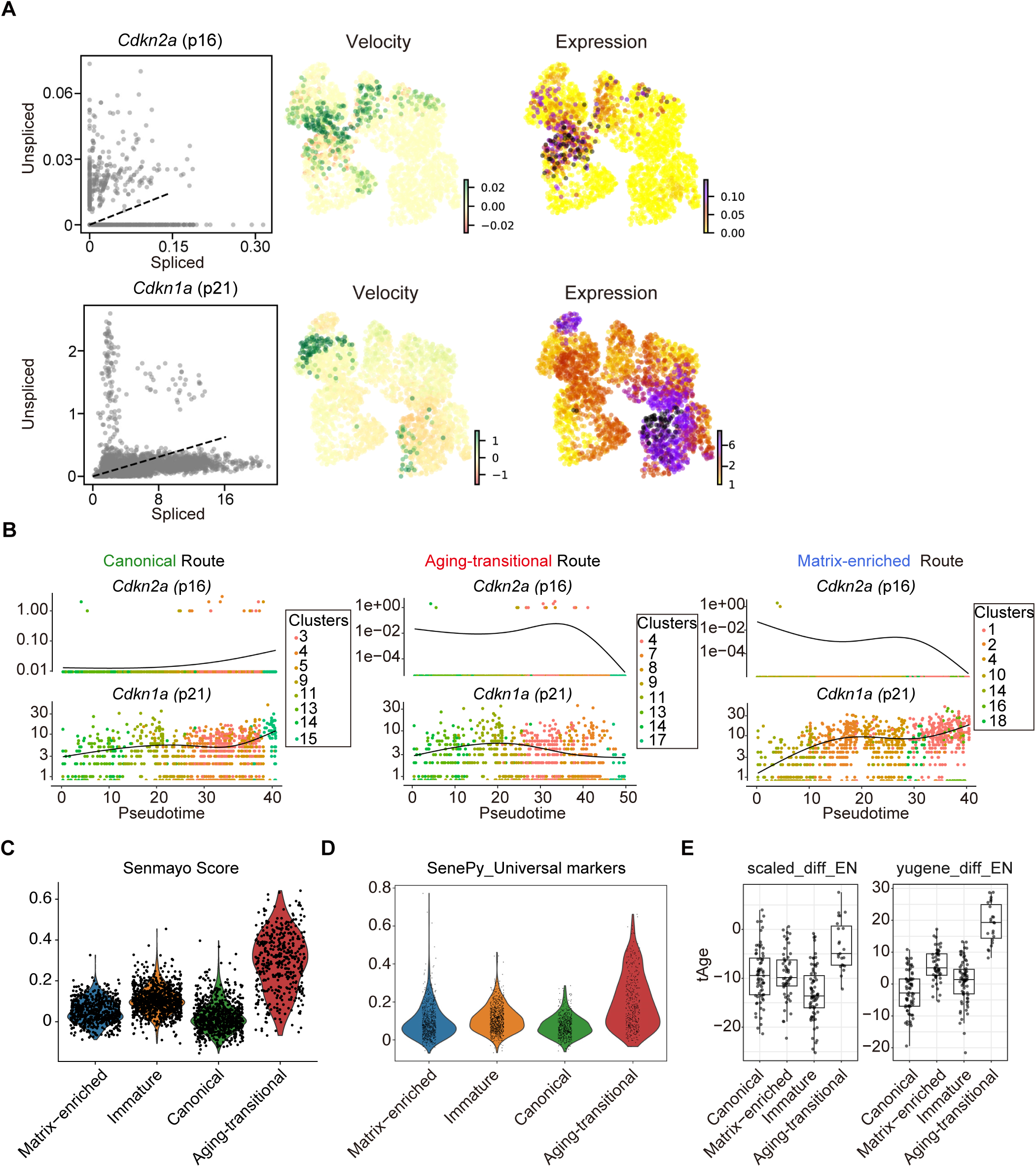
Osteocyte state transitions are early events that partially precede overt senescence. **(A)** RNA velocity and expression levels of *Cdkn2a* and *Cdkn1a* analyzed by scVelo. **(B)** Pseudotime analysis of *Cdkn2a* and *Cdkn1a* across distinct maturation routes using Monocle3. **(C)** Visualization of gene set scoring based on the SenMayo gene set. **(D)** Comprehensive evaluation of the senescence status of different osteocyte subpopulations using SenePy with universal markers. **(E)** Comprehensive evaluation of the aging status of different osteocyte subpopulations using tAge.

We then evaluated the functional contribution of senescent cells using systemic senolytic treatment with dasatinib and quercetin (D + Q) in middle-aged mice, a regimen reported to reduce senescent mesenchymal cells, including a fraction of osteocytes/late osteoblasts (Figs. 10A–C)(28). Despite the presence of senescence-associated gene expression in our dataset, D + Q treatment had limited effects on bone mass and remodeling parameters (Figs. 10D, E), in contrast to previous reports in aged models with more advanced senescent cell accumulation. Collectively, these results indicate that osteocyte state transitions precede overt senescence and cannot be fully explained by the accumulation of senescent osteocytes.

**Figure 10.**
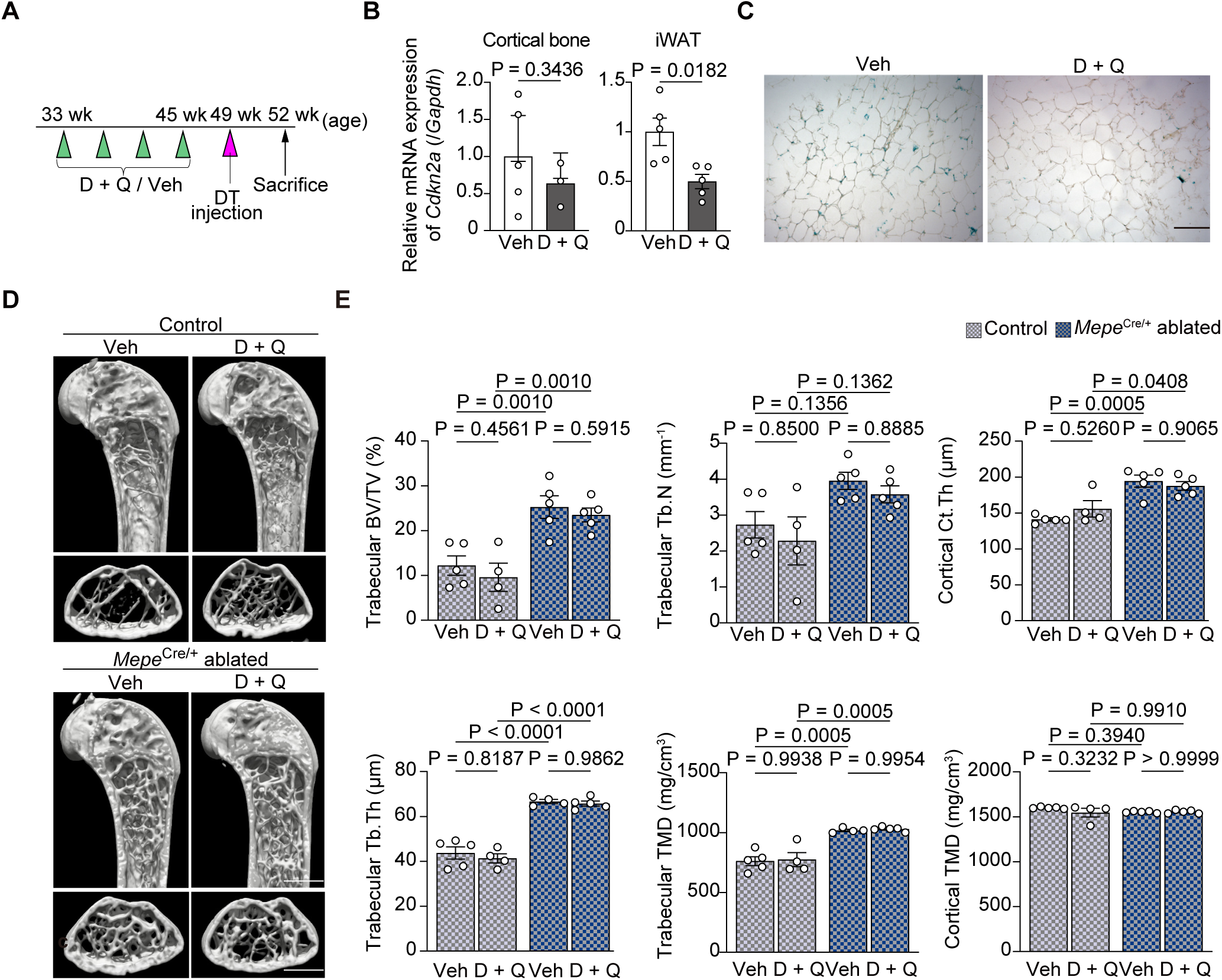
D + Q treatment does not recapitulate the increase in bone mass induced by osteocyte ablation in middle-aged mice. **(A)** Experimental schedule for the dasatinib/quercetin (D + Q) administration and ablation of *Mepe*-expressing osteocytes in 1-year-old mice. DT was administered 3 weeks before the analysis. D + Q was administered monthly starting 4 months before analysis. Veh, vehicle. **(B)** *Cdkn2a* expression in cortical bone and inguinal white adipose tissue (iWAT) from 1-year-old male *Mepe*^Cre/+^ *Rosa26*^LSL–DTR/LSL–DTR^ mice (n = 5 mice per group) 3 weeks after DT injection with or without D + Q treatment. **(C)** Representative senescence-associated β-galactosidase (SA-β-gal) staining image of iWAT from 1-year-old male *Mepe*^Cre/+^ *Rosa26*^LSL–DTR/LSL–DTR^ mice 3 weeks after DT injection with or without D + Q treatment. Scale bar,100 μm. **(D, E)** μCT analysis of distal femurs from 1-year-old male *Rosa26*^LSL–DTR/LSL–DTR^ and *Mepe*^Cre/+^ *Rosa26*^LSL–DTR/LSL–DTR^ mice (n = 3–5 mice per group) 3 weeks after DT injection with or without D + Q treatment. Scale bars, 1 mm. All data are presented as mean ± SEM. Statistical significance was determined by two-way ANOVA with Tukey’s test after excluding outliers via Grubbs’ test.

### Transcriptional and ligand–receptor analyses identify candidate regulators of osteocyte state transitions

Given that senescence alone does not fully account for the observed osteocyte state transition, we next explored potential upstream regulators by integrating ligand– receptor prediction with transcriptional regulatory network analyses. Using the NicheNet(29) package, we inferred ligand–receptor interactions based on receptor expression across osteocyte subpopulations, identifying several candidate signaling molecules, including *Tgfb1*, *Wnt1*, *Fgf2*, and BMP family members (Figs. S7–13). Distinct ligand profiles were associated with each subpopulation; extracellular matrix– related and remodeling-associated ligands were enriched in the aging-transitional subpopulation, whereas developmental and anabolic signaling molecules were more prominent in the matrix-enriched subpopulation.

In parallel, transcription factor activity was assessed using the pySCENIC(30) pipeline across both narrow and broad cis-regulatory regions (Fig. S14). Integration of pySCENIC predictions with NicheNet-inferred target gene sets allowed us to refine candidate regulators and identify overlapping transcriptional programs. This analysis highlighted several transcription factors associated with each state, including *Hmga2*, *Elk3*, *Sox9*, and *Hnf4a* in the matrix-enriched subpopulation, and *Cebpb*, *Nfkb1*, and *Ikzf2* in the aging-transitional subpopulation (Fig. 11). Together, these analyses identified distinct ligand–receptor interactions and transcription factor programs associated with each osteocyte subpopulation, indicating coordinated regulatory programs that may underlie osteocyte state transitions.

**Figure 11.**
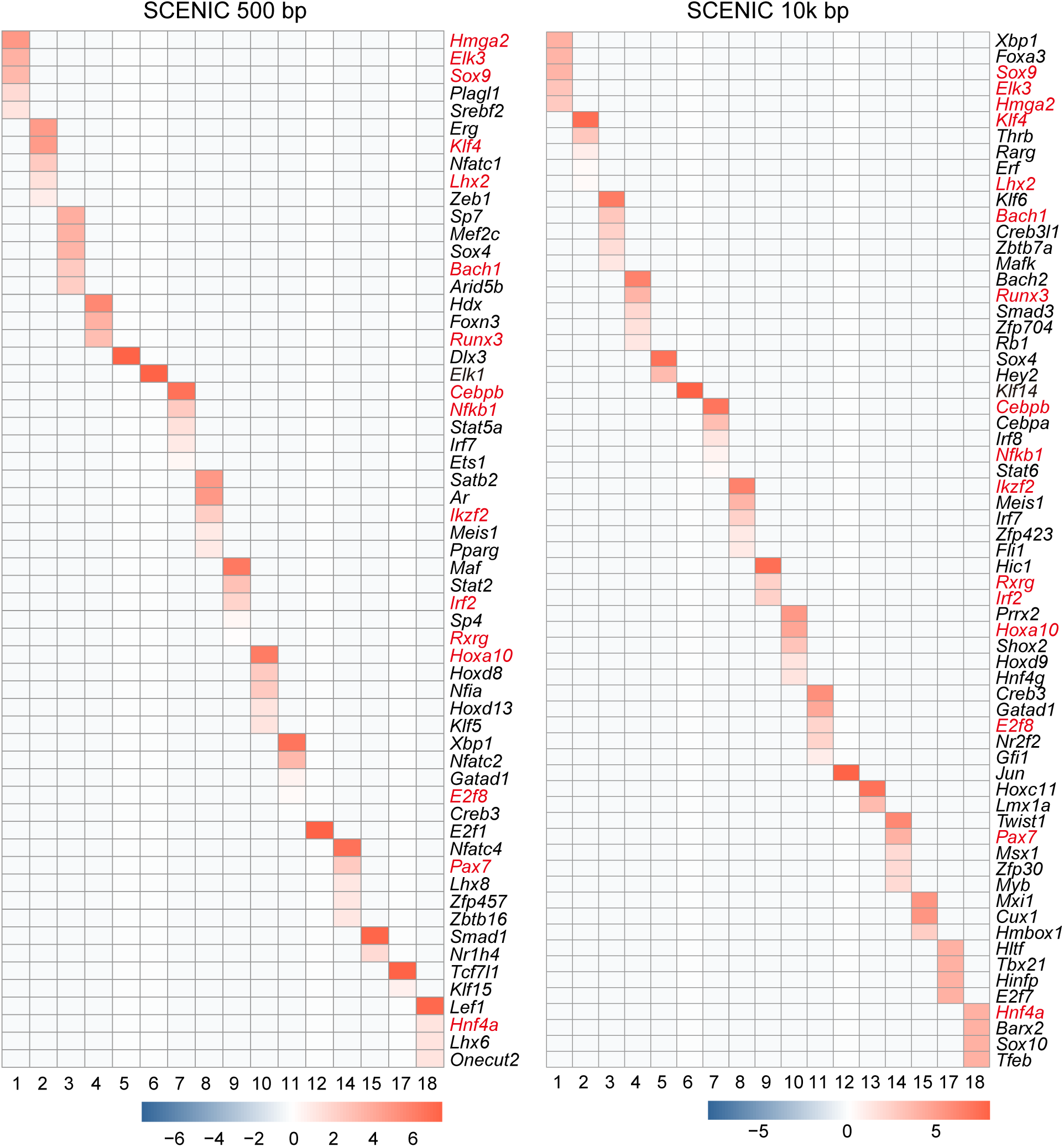
Transcriptional and ligand–receptor analyses identify candidate regulators of osteocyte state transitions. Heatmaps of candidate transcription factors weighted by the overlap between ligand target genes (from NicheNet) and regulons (from pySCENIC). Red highlights indicate transcription factors consistently identified across different pySCENIC threshold settings.

## Discussion

SIBLINGs (small integrin-binding ligand N-linked glycoproteins) are a family of proteins that regulate mineralization and phosphorus metabolism in hard tissues, and contain ASARM (acidic Ser/Asp-rich motif) peptides that can be released through proteolytic cleavage(31, 32). Among them, DMP1 and MEPE (matrix extracellular phosphoglycoprotein) are highly expressed within osteocyte lineage and interact with PHEX (phosphate regulating endopeptidase homolog, X-linked) to modulate FGF23 expression(33). In this study, we developed *Mepe*^Cre^, which utilizes the endogenous *Mepe* regulatory elements to achieve osteocyte-enriched Cre expression. Compared with *Dmp1*–Cre, which has been reported to exhibit off-target activity in non-skeletal tissues(19), *Mepe*^Cre^ demonstrated more restricted and stable expression in osteocytes, enabling more precise interrogation of osteocyte function *in vivo*. Previous studies(8) provide corroborative evidence for the utility of *Mepe*^Cre^, enabling us to revisit osteocyte function with greater specificity, hence offering a clearer framework for interpreting osteocyte-dependent phenotypes.

Osteocytes were once considered passive remnants of osteoblast differentiation, but are now recognized as key regulators of bone metabolism, orchestrating bone matrix remodeling, PTH-induced bone turnover, mechanical stress response, and the regulation of osteoblast and osteoclast activities(2, 4, 6). Nevertheless, their essential physiological roles remain incompletely defined. Our results indicate that osteocytes in young mice contribute to the suppression of bone formation under physiological conditions. Serum phosphorus concentrations remained unchanged in our study (Table S1). Although phosphate homeostasis can be mediated through multiple compensatory mechanisms, this finding, together with a previous study showing the regulation of bone formation by MEPE, PHEX, and ASARM peptides(34), suggests that the bone formation-suppressive function of osteocytes in young mice is unlikely to be mediated by the MEPE–PHEX– FGF23 or PHEX–FGF23 axis. The increase in bone formation observed following osteocyte ablation may therefore reflect the removal of osteocyte-derived inhibitory signals. One potential mechanism underlying this effect is the reduction of sclerostin levels(35–37), although additional regulatory pathways are likely involved given the pleiotropic functions of osteocytes. Taken together, these findings suggest that the suppression of bone formation represents a key physiological function of osteocytes in young mice, while leaving the precise molecular basis open for future investigation.

The link between osteocytes and aging has emerged as a major research focus, in line with accumulating clinical evidence implicating osteocyte alterations in age-related bone loss(38–40). It has been shown that 12–18 months of age represents a critical period for osteocyte senescence in mice, and that both systemic and osteocyte-targeted senolysis can increase bone mass, highlighting the importance of osteocyte senescence in late-life bone metabolism(14, 15, 27, 41, 42). In the present study, we identified a marked shift in osteocyte subpopulations during aging, characterized by a transition from the matrix-enriched to aging-transitional states. Notably, this shift was already evident at 12 months of age, suggesting that osteocyte remodeling begins relatively early in the aging process, prior to the establishment of overt senescence. Aging is a multifactorial process involving not only cellular senescence but also mitochondrial dysfunction, hormonal shifts, chronic inflammation, changes in cellular composition within tissues, and disrupted intercellular communication(43). Consistent with a previous study(44), our results show that bone loss begins between 6 and 12 months of age, prior to reaching an advanced age, a phenomenon similarly observed in humans(45). These observations together support the hypothesis that changes in osteocyte subpopulation composition may contribute to the initiation of bone loss during early-stage aging, preceding the accumulation of fully senescent cells. Mechanistically, these shifts likely involve coordinated changes in transcriptional programs and signaling pathways, as exemplified by a transition from cartilage-like to proinflammatory signatures, although the upstream drivers remain to be defined.

In this study, to achieve higher-resolution profiling of osteocytes, we performed scRNA-seq on bone cells sorted based on *Mepe*^Cre^-driven tdTomato fluorescence. Although this strategy facilitated a focused analysis of osteocyte subpopulations, it inherently precluded the investigation of broader intercellular interactions with neighboring cell types. Therefore, the upstream and downstream cues that govern the osteocyte state transitions identified here remain to be fully defined. Nevertheless, several lines of prior evidence are consistent with our observations, even if they do not directly explain the subpopulation transitions described in this study: newly formed osteocytes preferentially express *Tnfrsf11b* over *Tnfsf11*(46); osteocyte-derived RANKL plays a pivotal role in age-related bone loss and is upregulated in association with cellular senescence(47); and a C/EBP-binding intronic enhancer has been identified within the *Tnfsf11* gene locus(48). Another important consideration is that our analysis was conducted on 12-month-old mice, which are generally considered middle-aged rather than advanced-aged. This age window may be particularly informative for capturing a previously less-defined phase of osteocyte state remodeling that precedes the full manifestation of overt senescence-associated phenotypes. Experiments under conditions of advanced aging, in which p16-expressing senescent cells are more prevalent and other hallmarks of aging are more fully manifested, remain a subject for future investigation.

This study has several limitations. Although the *Mepe*^Cre^ model exhibits improved osteocyte specificity compared with *Dmp1*–Cre, minor recombination activity in late-stage osteoblasts/pre-osteocytes cannot be entirely excluded. Our conclusions are based on murine data and may not fully reflect human bone physiology. While our single- cell transcriptomic and metabolic analyses provided mechanistic clues, direct functional validation at the cellular level remains to be addressed in future studies. In addition, technical challenges in osteocyte isolation restricted the total cell yield available for single-cell analysis. Nevertheless, the use of fluorescence-based purification, together with consistent clustering and trajectory patterns across analyses, supports the robustness of the identified osteocyte subpopulations.

In summary, using the newly established *Mepe*^Cre^ model, we demonstrate that osteocytes suppress bone formation in young mice and undergo age-dependent functional changes. These changes reflect osteocyte state transitions characterized by coordinated alterations in signaling, transcriptional regulation, and metabolic profiles. Our results support a model in which osteocyte state transitions, rather than uniform osteocyte activity or senescence alone, govern bone remodeling across life stages.

## Methods

All experiments in this study were approved by the Institutional Animal Care and Use Committee/Genetically Modified Organisms Safety Committee/Research Safety Control Office of Institute of Science Tokyo.

### Mice, animal housing and basic animal experimental procedure

The generation of *Mepe*^Cre^ mice is described in Text S1(49). All mouse strains were maintained on a C57BL/6J background. Mice were housed by strain in a specific pathogen-free facility maintained at 23–25 °C and 40–70% humidity, under a 12-hour light/dark cycle, with ad libitum access to water and standard laboratory chow. Only male mice were used to avoid potential confounding effects of sex hormones and the estrous cycle on age-related skeletal remodeling. Therefore, the findings reported here are directly applicable to male mice, and future studies will be required to determine whether similar osteocyte state transitions occur in females.

For postnatal activation of *Rosa26*-iDTR, 7- or 49-week-old male mice received intraperitoneal injections of DT (50 µg/kg body weight/day) or PBS (vehicle), either once or as described in the figures and figure legends.

Age-matched littermates were used for all analysis. Mice received subcutaneous injections of calcein (16 mg/kg body weight) 7 days and 2 days (middle-aged) or 4 days and 1 day (young) prior to sacrifice. After the mice were euthanized via CO_2_ inhalation, blood and bone samples were collected for downstream analyses. The number of mice used and the experimental schedules are described in the respective figures or figure legends.

### μCT (micro-computed tomography) Analysis

For three-dimensional μCT analysis, right femurs were fixed in 70% ethanol for at least 1 week. μCT scanning was performed using a ScanXmate-A100S scanner (Comscantechno). Three-dimensional microstructural images were reconstructed, and structural parameters were calculated using TRI/3D-BON software(50).

### Histological analysis

For bone histomorphometric analysis, undecalcified right tibiae were embedded in glycol methacrylate, sectioned at 5 μm, and stained with toluidine blue, tartrate-resistant acid phosphatase (TRAP), or using the Villanueva method. Parameters for osteoblasts, osteocytes, and osteoclasts in the secondary trabecular bone were assessed in microscopic fields from two sections per mouse by systematically moving the field of view across the x- and y-axes.

For hematoxylin and eosin (HE) staining, immunohistochemistry, and osteocyte apoptosis assessment, left tibiae were fixed in 4% paraformaldehyde (PFA)/PBS for 24 h at 4 °C, decalcified in EDT-X for 2–3 weeks, dehydrated, and embedded in paraffin. Soft tissues were processed similarly, except for the decalcification step. HE and immunohistochemical staining (Primary antibody: Goat anti-human HB-EGF, 1:40; Secondary antibody: Histofine Simple Stain Mouse MAX PO (G), ready to use) were performed as previously described(4, 51). Osteocyte apoptosis was evaluated by both HE staining and the DeadEnd™ Fluorometric TUNEL System, with DAPI counterstaining using VECTASHIELD Mounting Medium.

For cryostat sectioning in reporter mouse analysis, mice were administered calcein twice prior to being euthanized and perfused with 4% PFA/PBS. The left femurs were then harvested and post-fixed in 4% PFA/PBS at 4 °C for 1 h. Other tissues were fixed in 4% PFA/PBS for 12 h and subsequently immersed in 30% sucrose/PBS until fully submerged. Undecalcified sections (8 μm) were prepared using Kawamoto’s method with a cryostat (Leica Biosystems). After dehydration, sections were either mounted in VECTASHIELD or stained using the Senescence β-Galactosidase Staining Kit.

Images were captured using an all-in-one fluorescence microscope (BZ-X700, KEYENCE). Bone histomorphometric analyses were performed using WinROOF 2013 software, and other image analyses were performed using the BZ-X Analyzer. For quantification of tdTomato-positive cells, femoral sections were anatomically divided into predefined regions of interest (ROIs), defined as cortical bone, trabecular bone, and the articular region. tdTomato-positive cells were counted only when tdTomato fluorescence was clearly associated with a DAPI-positive nucleus, thereby minimizing false-positive signals derived from nonspecific fluorescence or tissue autofluorescence. Calcein labeling was used to identify active mineralization fronts and to assist with anatomical orientation. For each mouse, at least four non-consecutive sections were analyzed.

### Serum analysis

After blood collection, samples were left at room temperature for 30–60 minutes in microtubes, centrifuged at 10,000 rpm for 10 minutes, and the serum was collected and stored at –80 °C. Serum biochemical profiling was performed by Oriental Yeast. Concentrations of FGF23 and sclerostin were measured using the Mouse & Rat SOST/Sclerostin ELISA Kit and the FGF-23 ELISA Kit, respectively, according to the manufacturers’ instructions.

### Quantitative RT-PCR

Adipose tissue was homogenized in TRI Reagent, and total RNA was isolated according to the manufacturer’s instructions. First-strand cDNA was synthesized using the PCR Thermal Cycler Dice Touch (TaKaRa Bio), employing the ReverTra Ace qPCR RT Master Mix with gDNA Remover. Quantitative PCR was performed using the CFX384 Touch Real-Time PCR Detection System and CFX Manager software with SYBR™ Green Realtime PCR Master Mix. Relative gene expression levels were calculated using the ΔΔCt method and normalized to the internal reference gene *Gapdh*. Primer sequences used in this study are as follows: *Gapdh*, F: AACTTTGGCATTGTGGAAGG, R: GGATGCAGGGATGATGTTCT; *Cdkn2a,* F: CCCAACGCCCCGAACT, R: GCAGAAGAGCTGCTACGTGAA; *Mepe*, F: GATGCAGGCTGTGTCTGTTG, R: TGTCTTCATTCGGCATTGG; *Sost*, F: TCCTGAGAACAACCAGACCA, R: GCAGCTGTACTCGGACACAT; *Pdpn*, F: GCAGGGGATGAAACGCAGA, R: TAGCTCTTTAGGGCGAGAACC; *Dmp1*, F: CCCAGAGGGACAGGCAAATA, R: TCCTCCCCACTGTCCTTCTT.

### Senescent cell deletion

To induce senolysis, 32-week-old mice were administered Dasatinib (D; 5mg/kg body weight) and Quercetin (Q; 50mg/kg body weight) via oral gavage once a month for 4 months. Control mice received vehicle (10% polyethylene glycol 4000 solution) alone.

### Flow cytometry analysis of immune cells

For flow cytometry analysis of the spleen and thymus, single-cell suspensions were prepared by mechanically dissociating the tissues through a 60 µm cell strainer and flushing with 2% FBS/PBS, centrifuged, incubated with TruStain FcX™ to block Fc receptors, and stained with specific antibodies for 30 min on ice. After 3 washes with 2% FBS/PBS, cells were stained with 7-AAD and analyzed using a FACS Aria III and associated software (BD Biosciences). Antibodies conjugated with fluorescein isothiocyanate (FITC), phycoerythrin (PE), allophycocyanin (APC), PE-Cy7, and APC- Cy7 were used for surface and intracellular staining. The following antibodies were obtained from BioLegend: anti-mouse CD3 (1:500), CD4 (1:500), CD8a (1:500), CD11b (1:500), CD19 (1:500), CD45R (1:500). A detailed gating strategy is provided in Supplementary Information (Figs. S15A and S15B).

### Osteocyte-rich cell isolation and data acquisition for scRNA-seq

Osteocyte-rich cells were harvested as previously described(52). Briefly, 12- and 52- week-old *Mepe*^Cre/+^ *Rosa26*^CAG–tdTomato/+^ mice (n = 3 per group; samples from 3 mice were pooled) were euthanized, and tibiae, femurs, and humeri were collected. Soft tissues were removed by scraping, and the epiphyses were excised. Bone marrow was flushed out with PBS. Bones were then cut into ∼2 mm fragments and subjected to eight serial enzymatic digestions as described previously. Freshly isolated cells from fractions 4–8 were collected, resuspended in 2% FBS/PBS containing RNase inhibitor, and filtered through a 60 µm cell strainer.

CD45-positive cells were depleted using the EasySep Mouse CD45 Positive Selection Kit, following the manufacturer’s instructions, to enrich for mesenchymal cells. After DAPI staining, live, single, tdTomato-positive cells were sorted using a FACS Aria III (BD Biosciences) according to the gating strategy described (Fig. S15C).

Sorted cells were encapsulated into emulsion droplets using the Chromium Controller (10x Genomics). For each group, over 5,000 cells were loaded to obtain a single library using the Chromium Next GEM Single Cell 3′ v3 chemistry. Libraries were barcoded, purified, and sequenced in a 2 × 150 bp paired-end format on a HiSeq platform (Illumina) at a depth of approximately 400 million reads.

### Data analysis for scRNA-seq

Cell Ranger was used to build a custom reference using the *Mus musculus* GRCm39 genome and GRCm39.113 gene annotation (both obtained from Ensembl), demultiplex and align reads, and generate the possorted BAM file and unfiltered feature-barcode matrices. Velocyto(26) was used to generate loom files containing spliced/unspliced count data based on the same GRCm39.113 annotation and the UCSC mm39 repeat masker file.

Unfiltered matrices from two experiments were imported into the Seurat pipeline(53). Cell cycle scores (S and G2/M phases), mitochondrial gene percentage, hemoglobin gene percentage, and log10GenesPerUMI were calculated. Filtering criteria included: (1) genes expressed in >3 cells; (2) cells with >200 genes detected; (3) lineage marker gene expression (e.g., *Ptprc*, *Cd3e*, *Cd79b*, *Cd79a*, *Itgax*, etc.) <3; (4) nFeature_RNA <6000; (5) mitochondrial gene percentage <20%; (6) hemoglobin gene percentage <5%; and (7) log10GenesPerUMI >0.75. Mitochondrial, hemoglobin, and ribosomal genes were excluded prior to normalization and scaling, which included cell cycle and technical covariates.

After preprocessing, datasets from 12- and 52-week-old mice were integrated using HarmonyIntegration option, following feature selection via FindVariableFeatures (method = "vst", nfeatures = 2000, mean.cutoff = c(0, 10), dispersion.cutoff = c(0, Inf)) and principal components analysis (PCA). Principal components (PCs 1–27) were used for graph-based clustering and UMAP for dimensionality reduction. Cluster-specific marker genes were identified via FindMarkers, and the top 10 genes per cluster were used to define the cell type and exclude hematopoietic and endothelial-like clusters. Gene set scoring based on the osteocyte signatures(8) was used for the subclustering of osteocyte-lineage cells. The final Seurat object ready for analysis (“Exp. Seurat object”) was reprocessed from FindVariableFeatures to FindMarkers as above, using PCs 1–23 for downstream analysis.

For differentiation potential, CytoTRACEv2(54) was used to calculate differentiation scores. Metadata, UMAP coordinates, and cluster identities were exported and merged with loom files via Scanpy(55) and loompy for RNA velocity analysis using scVelo(56), following the authors’ recommended procedure. Monocle3(57) was used for pseudotime analysis, with root cell assignment guided by integrating CytoTRACEv2 and scVelo results.

For pathway enrichment, differentially expressed genes were recalculated for distinct osteocyte subtypes using FindMarkers, and GO analysis was performed using clusterProfiler(58) with the following parameters: pAdjustMethod = "BH", pvalueCutoff = 0.05, and qvalueCutoff = 0.05.

For metabolic profiles assessment, scFEA(59) was applied following the developers’ tutorial (reference file: cmMat_complete_mouse_c70_m168). The resulting flux matrix was merged into the Seurat object and visualized by t-SNE and heatmaps.

For brief evaluation of senescence signatures, the AddModuleScore function in Seurat were used for quantification with the SenMayo gene set(27). Meanwhile, the senePy(60) and tAge(61) were used for more comprehensive evaluation of senescence and aging status.

To evaluate the ligand–receptor interactions necessary for osteocytes, NicheNet(29) were used. Cluster-specific markers were identified via FindAllMarkers, and average expression was calculated. Genes expressed in >10% of cells were used as background. Each cluster was set as a receiver, and ligands/receptors were filtered by intersection with the NicheNet prior network. Ligand activity was predicted using the provided ligand- target matrix, and the top 30 ligands were selected based on corrected AUPR scores (reference files: ligand_target_matrix_nsga2r_final_mouse, lr_network_mouse_21122021, and weighted_networks_nsga2r_final_mouse.rds).

To assess transcription factor (TF) regulatory activity, pySCENIC(62) was applied to the filtered gene expression matrix extracted from the Exp. Seurat object, following the authors’ tutorial. The following reference files were used: mm10_10kbp_up_10kbp_down_full_tx_v10_clust.genes_vs_motifs.rankings.feather;m m10_500bp_up_100bp_down_full_tx_v10_clust.genes_vs_motifs.rankings.feather;moti fs-v10nr_clust-nr.mgi-m0.001-o0.0.tbl; allTFs_mm.txt. Regulon activity inferred by pySCENIC was post-processed and visualized in R using AUCell scores stored in a loom file. The regulon–gene incidence matrix and AUC matrix were extracted using the SCopeLoomR package. Cell identities were assigned based on Seurat cluster annotations. Regulon specificity scores (RSS) and Z-scores were computed to quantify cluster-specific regulon activity. For each cluster, the top 5–20 regulons were ranked by RSS, and the five most specific regulons were visualized.

To identify TFs whose regulon targets overlapped with predicted ligand-responsive genes, pySCENIC and NicheNet results were integrated as follows: (1) TF–target gene relationships were obtained from pySCENIC and formatted into a cluster-wise regulon table; (2) predicted ligand target genes from NicheNet were retrieved for each cluster; (3) overlaps between pySCENIC-defined regulon targets and NicheNet-predicted targets were quantified for each TF in each cluster. The TFs showing the highest overlaps were selected and visualized to highlight convergence between ligand-induced signaling and TF-regulated transcriptional programs.

### Statistical Analysis

Each experiment included multiple biological replicates (individual mice). All data are presented as the mean ± standard error of the mean (SEM) values of independent replicates. Sample sizes are indicated in the figure or corresponding figure legends and were based on previously published studies; no statistical method was used to predetermine sample size. Outliers that may be related to technical error were identified using Grubbs’ test (*α* = 0.05) and excluded. The investigators were not blinded to allocation during the experiments and outcome assessment. For comparisons between two groups, two-tailed unpaired *t* test with Welch’s correction was used. For multiple group comparisons, two-way ANOVA with Šídák’s or Tukey’s multiple-comparison test was used. *P* < 0.05 was considered statistically significant (\**P* < 0.05; \*\**P* < 0.01; \*\*\**P* < 0.001; \*\*\*\**P* < 0.0001). All statistical analysis was performed with Prism 10.

## Key resources table

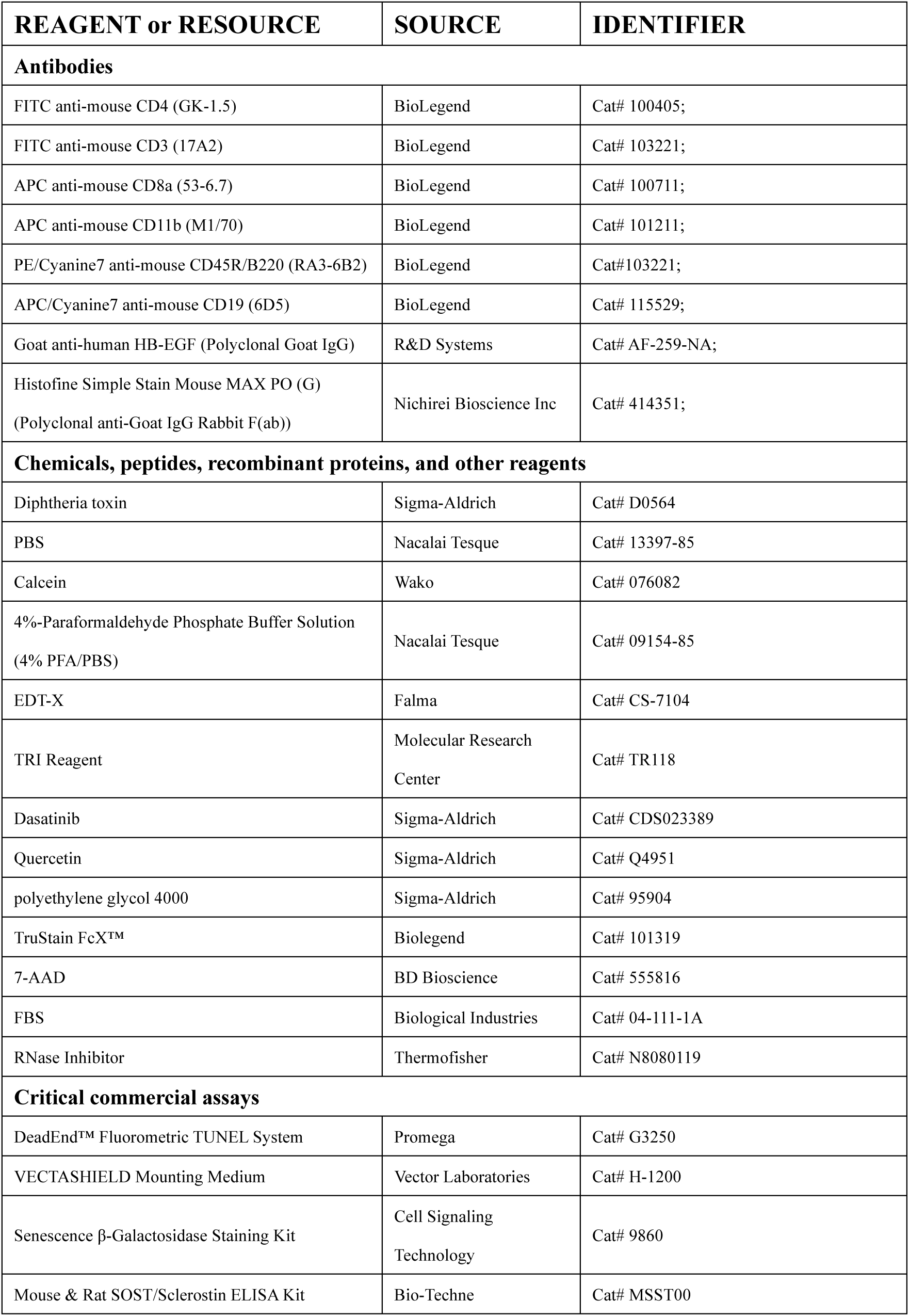
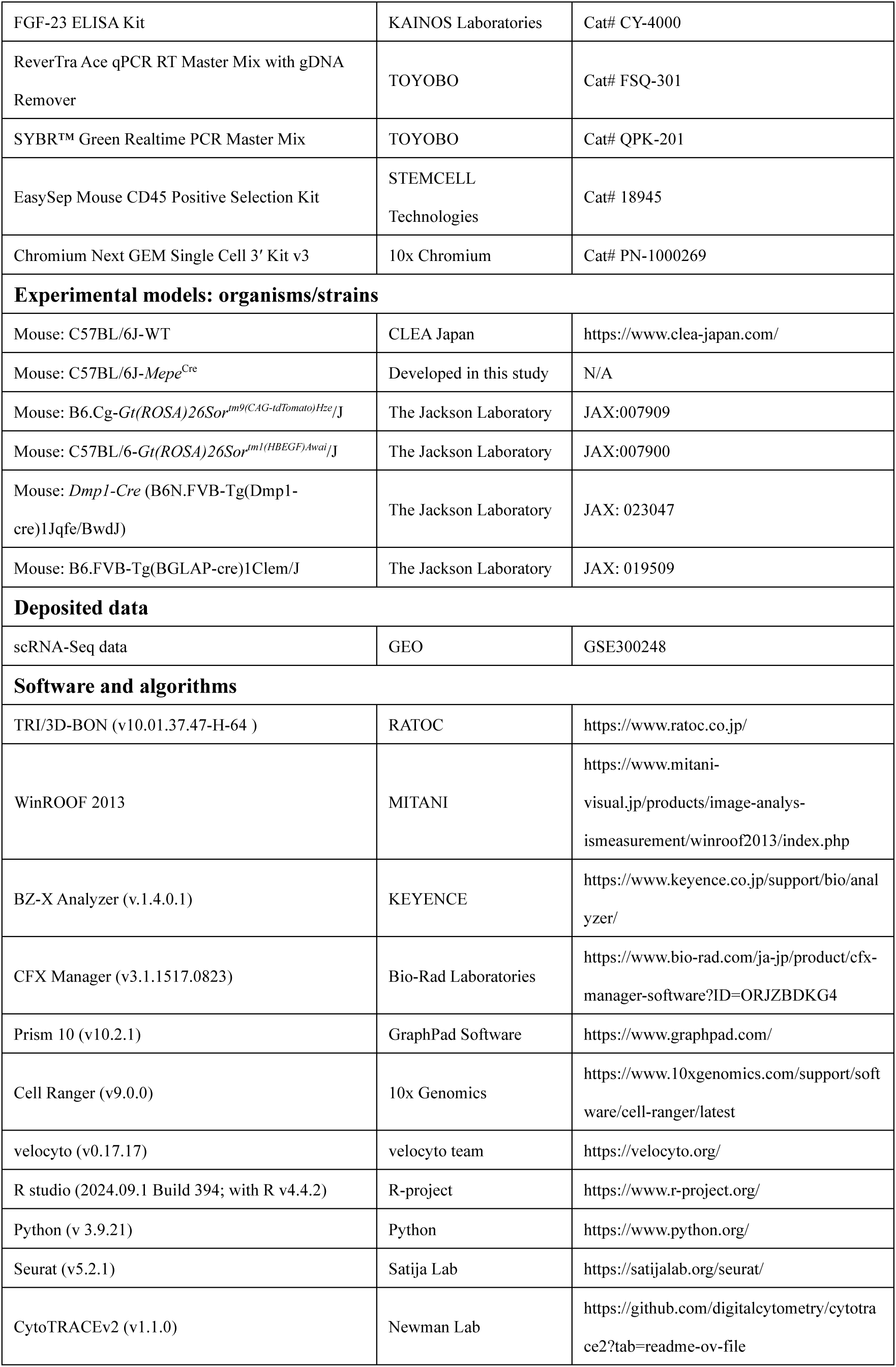
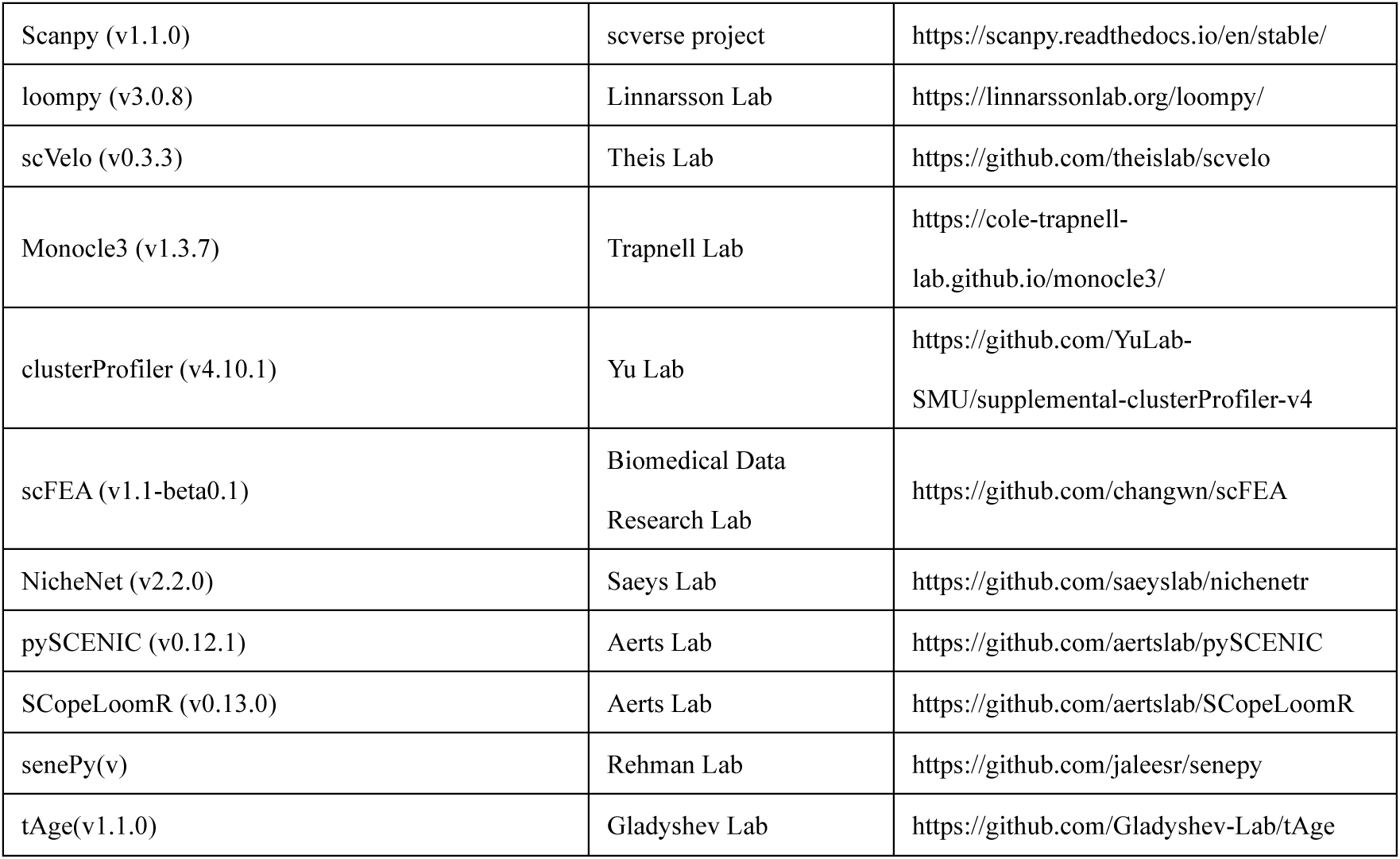

## Data Availability

The single-cell RNA sequencing data generated in this study have been deposited in GEO database under accession number GSE. Additional data are available from the corresponding author upon reasonable request. Source data for bar plots will be supplemented after the publishment.

## Author Contributions

R.D., A.L., M.H., and T.N. conceived and designed the research. R.D., A.L. and M.H. performed the research and acquired the data. R.D., A.L., M.H., and C.W. analyzed and interpreted the experiment data. A.L. and R.D. prepared the manuscript and figure with support from M.H., H.T., M.S., and T.N. H.A and T.N designed and generated the *Mepe*^Cre^ transgenic mice. The order of the co-first authors was determined solely based on the amount of experimental work performed by each author. All authors were involved in drafting and revising the manuscript.

## Acknowledgements

We thank K. Yusoon, M. Inoue, Y. Yamashita, T. Gondo, B Li and S. Kato for helpful discussions and technical assistance.

