## Supplemental Material for "Osteocyte State Transitions Modulate Bone Remodeling During Early Skeletal Aging"

**This PDF file includes:**

Figures S1 to S15
Table S1
Text S1

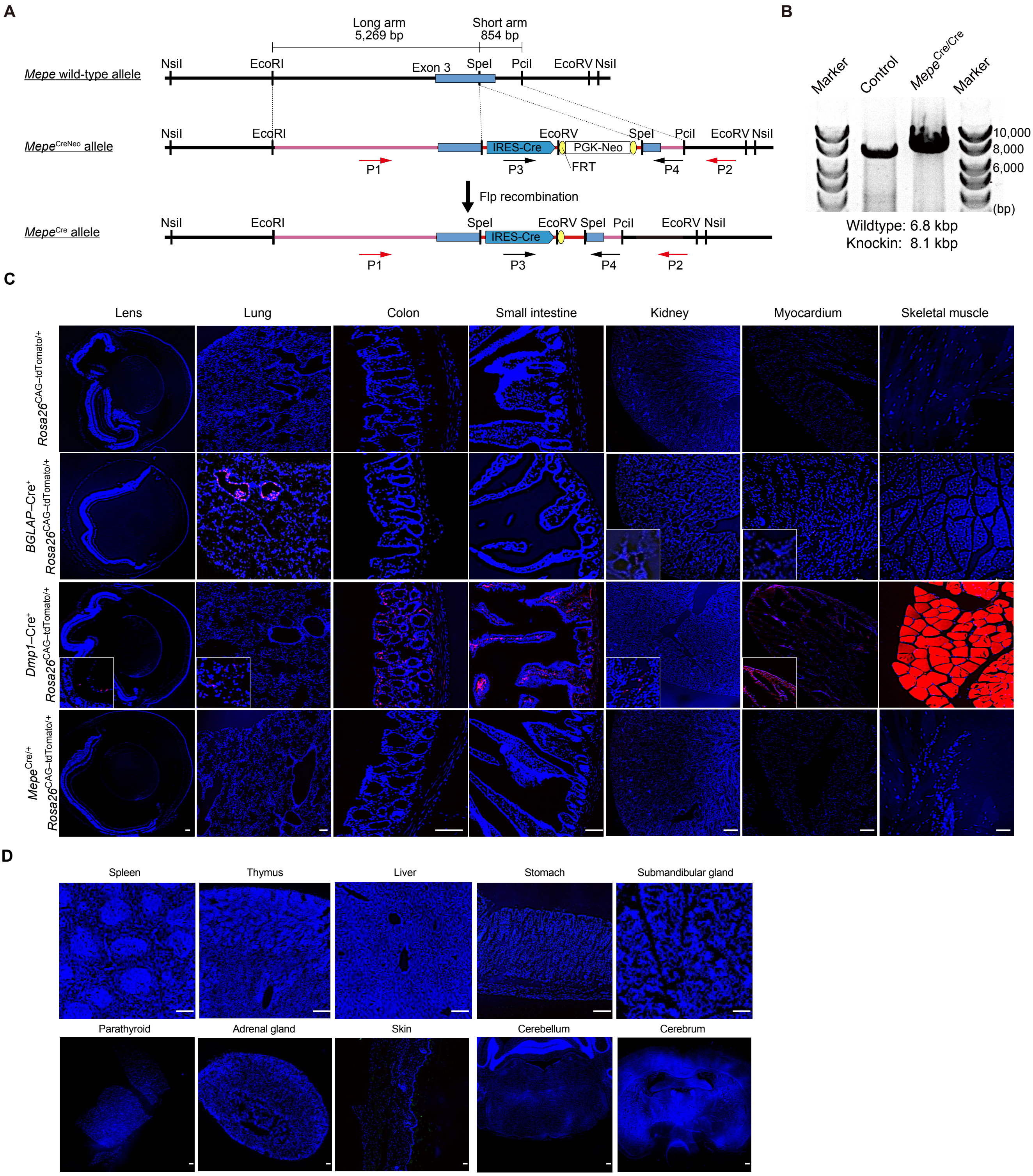


**Figure S1 Osteocyte-enriched Cre recombinase activity in *Mepe*$^{Cre/+}$ mice.**

**(A)** Targeting strategy to generate *Mepe-IRES*-Cre knock-in mice. The genomic structure of the wild-type *Mepe* gene, the targeting vector, and the targeted alleles are indicated. P1, forward long PCR primer; P2, reverse long PCR primer; P3, forward genotyping primer; P4, reverse genotyping primer. Specific sequences of the primers are described in Supplementary Text 1. **(B)** PCR analysis of tail genomic DNA using the P1/P2 primer pairs shown in Fig 1A. (**C)** Representative images of frozen sections of the lens, lung, colon, small intestine, kidney, myocardium and skeletal muscle from *Rosa26*$^{CAG–tdTomato/+}$, *BGLAP*–Cre$^{+}$ *Rosa26*$^{CAG–tdTomato/+}$, *Dmp1*–Cre$^{+}$ *Rosa26*$^{CAG–tdTomato/+}$, and *Mepe*$^{Cre/+}$ *Rosa26*$^{CAG–tdTomato/+}$ mice. Blue, DAPI. Scale bars, 100 μm. A magnified view of the positive cells is shown in the lower left corner of some images for clarity. (**D)** Representative images of frozen sections of the spleen, thymus, liver, stomach, submandibular gland, parathyroid, adrenal gland, skin, cerebellum, cerebrum from *Mepe*$^{Cre/+}$ *Rosa26*$^{CAG–tdTomato/+}$ mice. Blue, DAPI. Scale bars, 100 μ m.

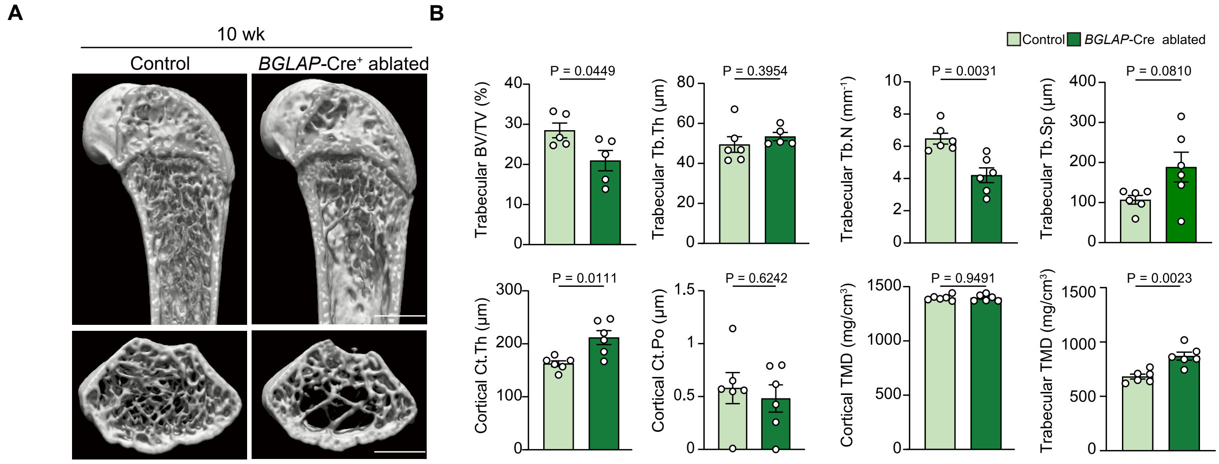


**Figure S2 *BGLAP*–Cre-driven ablation reduces the trabecular but increases cortical bone mass.**

**(A**, **B)** µCTanalysis of distal femurs from 10-week-old male *Rosa26*$^{LSL–DTR/LSL–DTR}$ and *BGLAP*-Cre$^{+}$ *Rosa26*$^{LSL–DTR/LSL–DTR}$ mice at 3 weeks after DT injection (biological replicate = 5–6). Scale bars, 1 mm.

All data are presented as mean ± SEM. Statistical significance was determined by Welch’ s t-test after excluding outliers via Grubbs’ method.

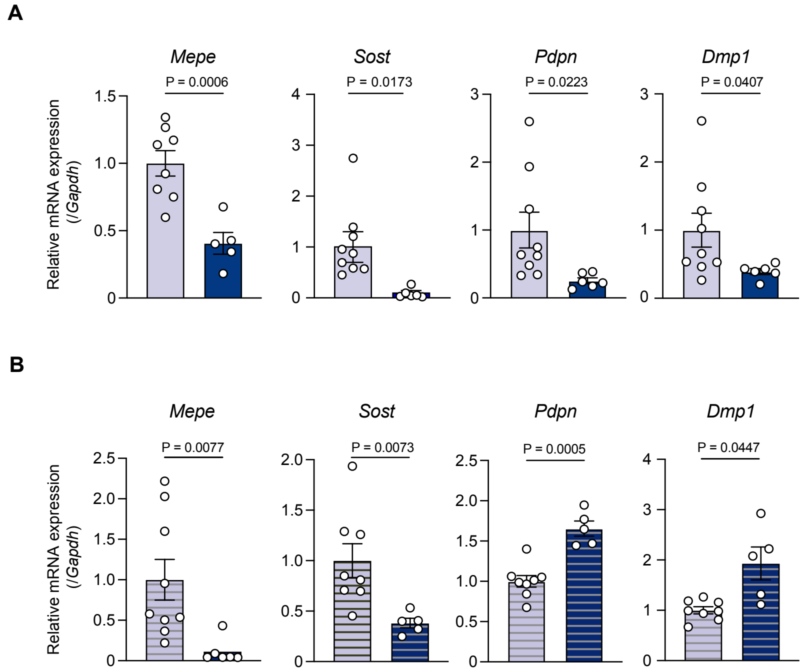


**Figure S3 Osteocyte-related gene expression in cortical bone following different ablation strategies after aging.**

**(A, B)** *Mepe, Sost, Pdpn, Dmp1* expression in bone and the serum sclerostin concentrations of 52-week-old male *Rosa26*$^{LSL–DTR/LSL–DTR}$ and *Mepe*$^{Cre/+}$ *Rosa26*$^{LSL–DTR/LSL–DTR}$ mice that received a single DT injection (A ;biological replicate = 5–8) or continuous DT injections (B; biological replicate = 6–9). All data are presented as mean ± SEM. Statistical significance was determined by Welch’ s t-test after excluding outliers via Grubbs’ method.

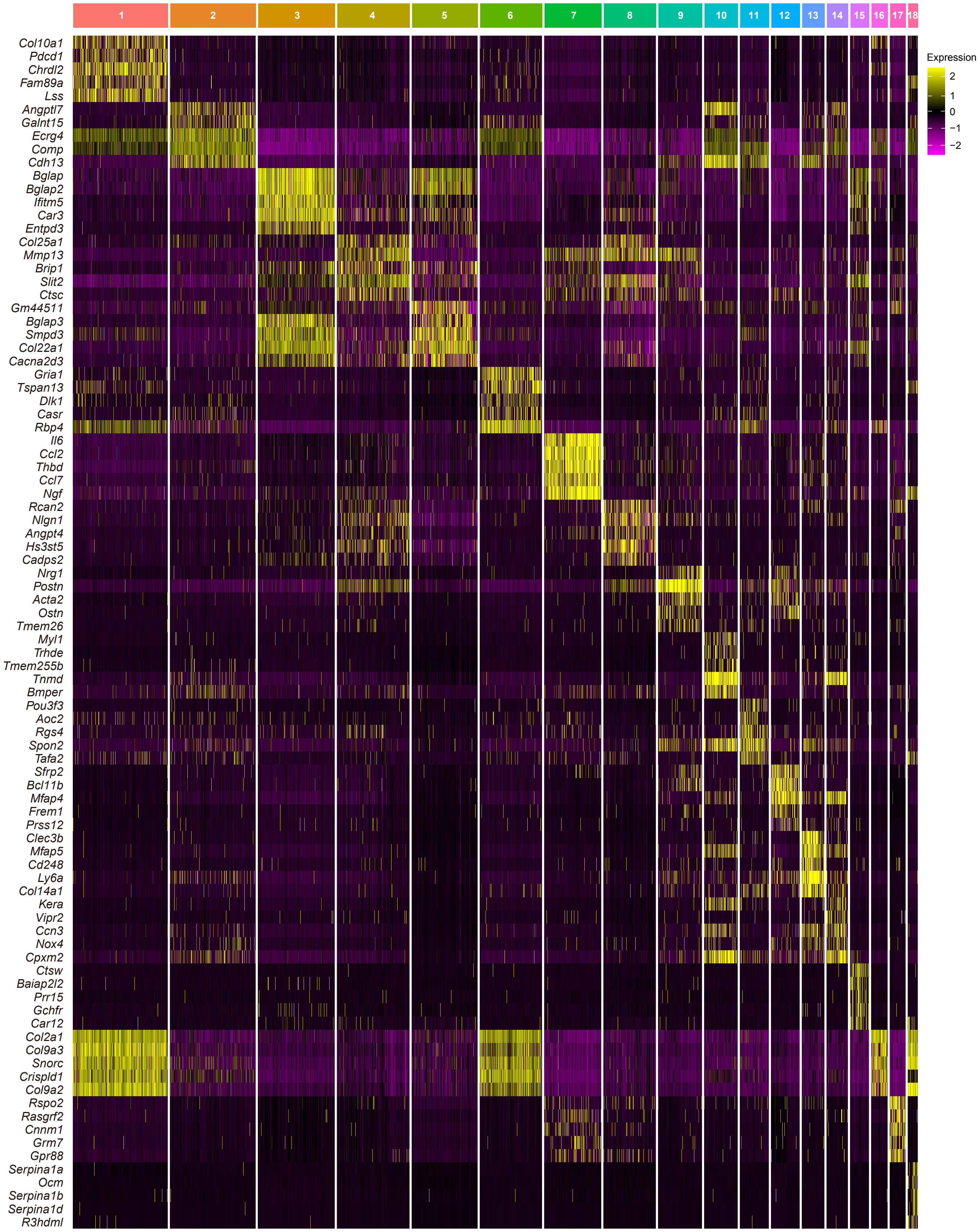


**Figure S4 Heatmap of markers for distinct osteocyte clusters.**

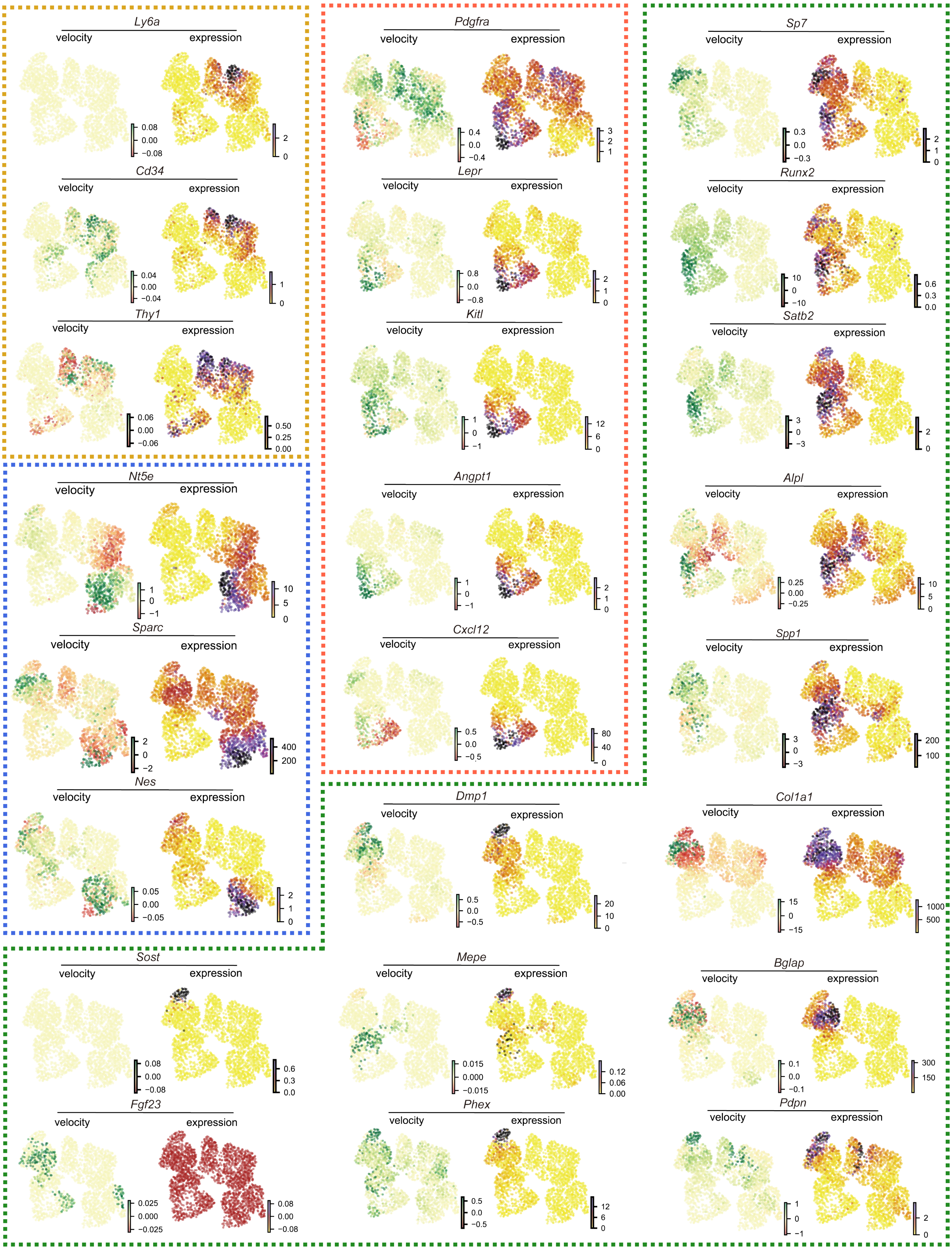


**Figure S5 RNA velocity and expression levels of osteolineage-related markers in *Mepe*$^{Cre/+}$-sorted osteocytes.**
Different outline colors indicate genes with enriched expression in distinct osteocyte subtypes. Yellow, Immature osteocytes; red, aging-transitional osteocytes; blue, matrix-enriched osteocytes; green, canonical osteocytes.

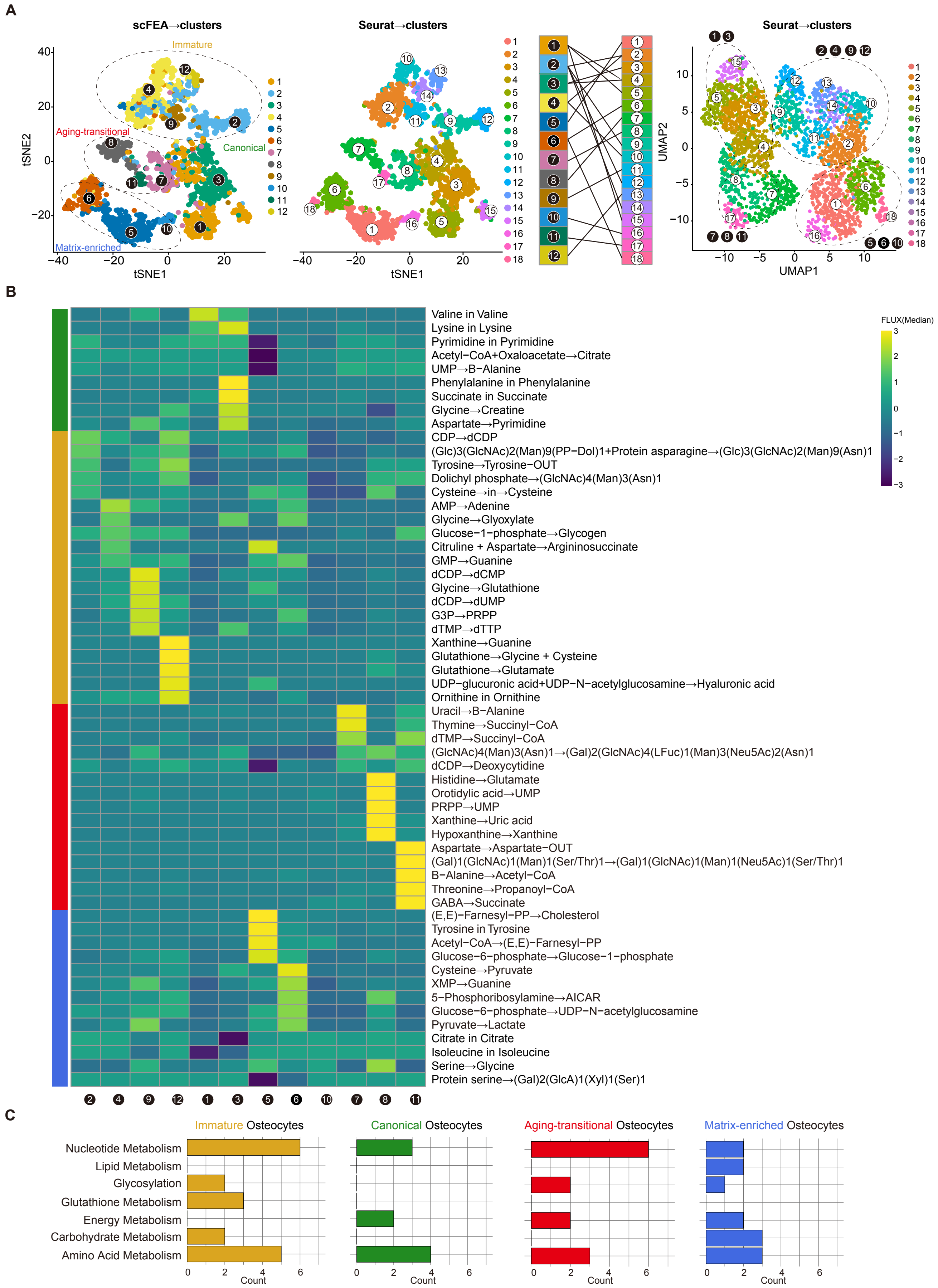


**Figure S6 Distinct metabolic profiles among *Mepe*$^{Cre/+}$-sorted osteocyte subtypes.**
**(A)** Metabolic profiles calculated by scFEA and visualized using tSNE. Seurat clusters in the prior UMAP are simultaneously plotted on tSNE to better visualize their relationships. **(B)** Heatmap of highly activated metabolic module. The top 5 modules in each scFEA clusters were shown according to Z-score. **(C)** Categorization of the module activated in distinct osteocyte subpopulations.

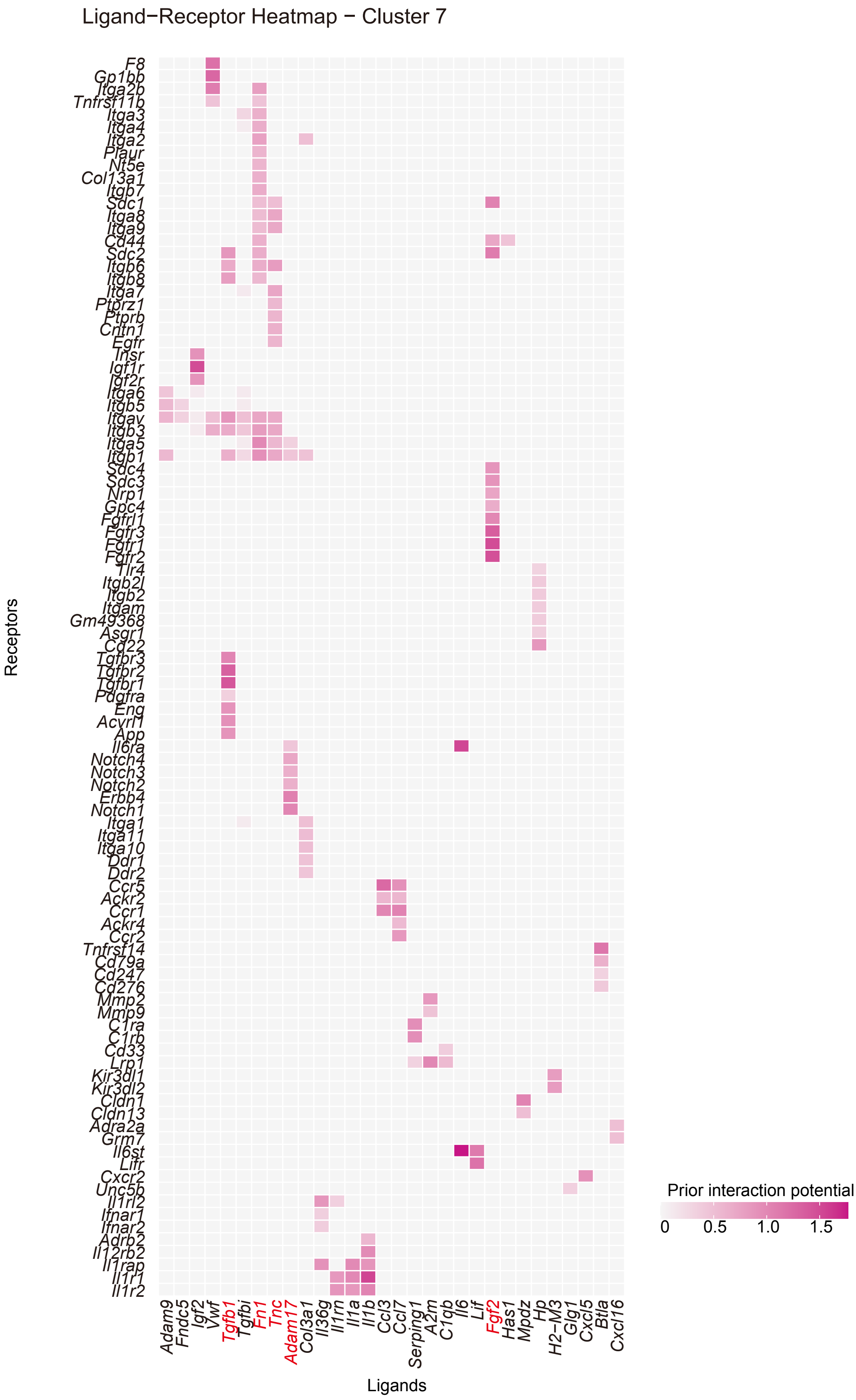


**Figure S7 Predicted signaling in the aging-transitional subpopulation (Seurat cluster 7) in *Mepe*$^{Cre/+}$-sorted osteocytes.**
Heatmap of predicted ligands (signal) and receptors activated in the aging-transitional cluster (cluster 7) are shown. Ligands highlighted in red correspond to the top five receptors that express more numbers of receptors in the clusters.

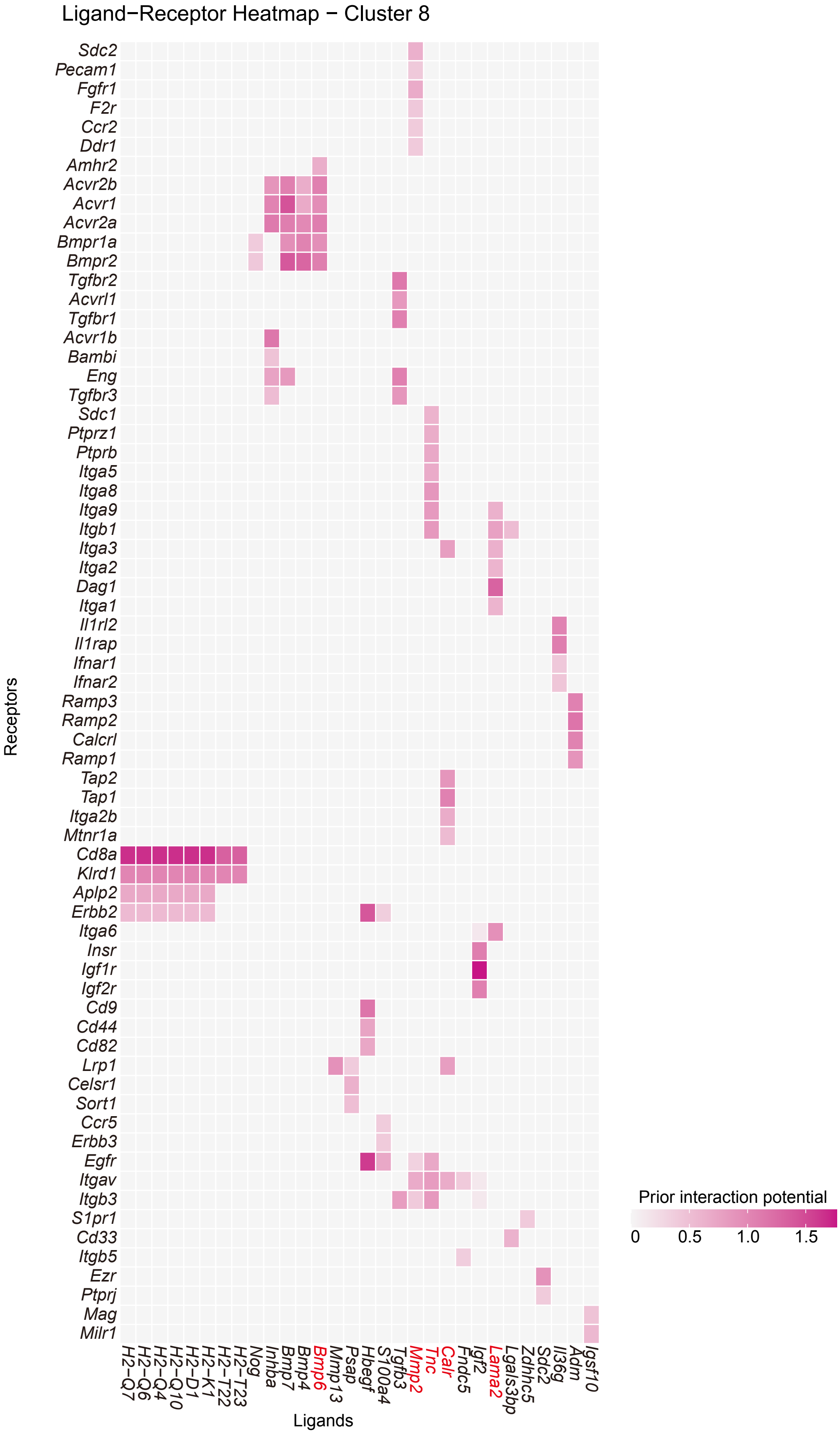


**Figure S8 Predicted signaling in the aging-transitional subpopulation (Seurat cluster 8) in *Mepe*$^{Cre/+}$-sorted osteocytes.** Heatmap of predicted ligands (signal) and receptors activated in the aging-transitional cluster (cluster 8) are shown. Ligands highlighted in red correspond to the top five receptors that express more numbers of receptors in the clusters.

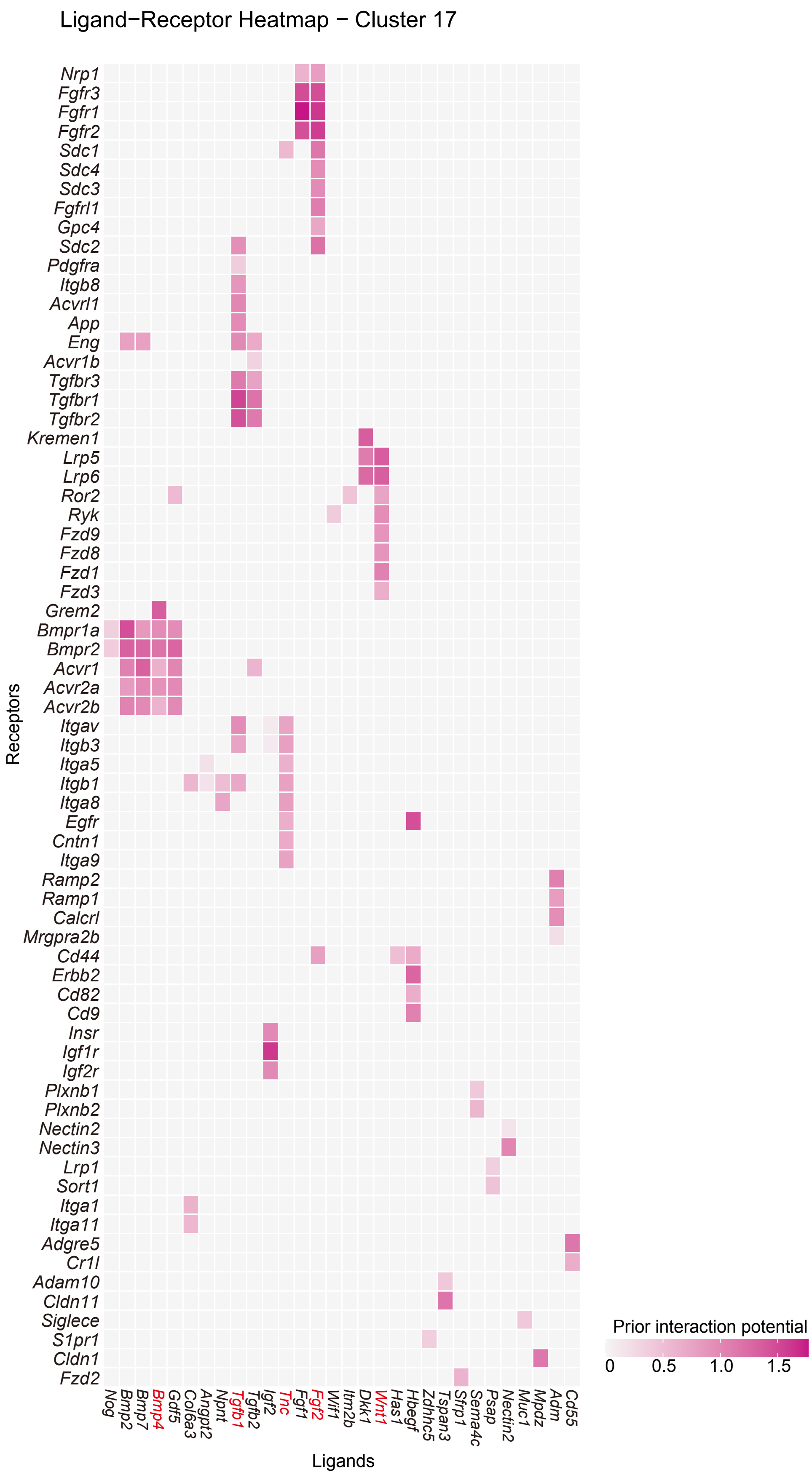


**Figure S9 Predicted signaling in the aging-transitional subpopulation (Seurat cluster 17) in *Mepe*$^{Cre/+}$-sorted osteocytes.**
Heatmap of predicted ligands (signal) and receptors activated in the aging-transitional (cluster 17) are shown. Ligands highlighted in red correspond to the top five receptors that express more numbers of receptors in the clusters.

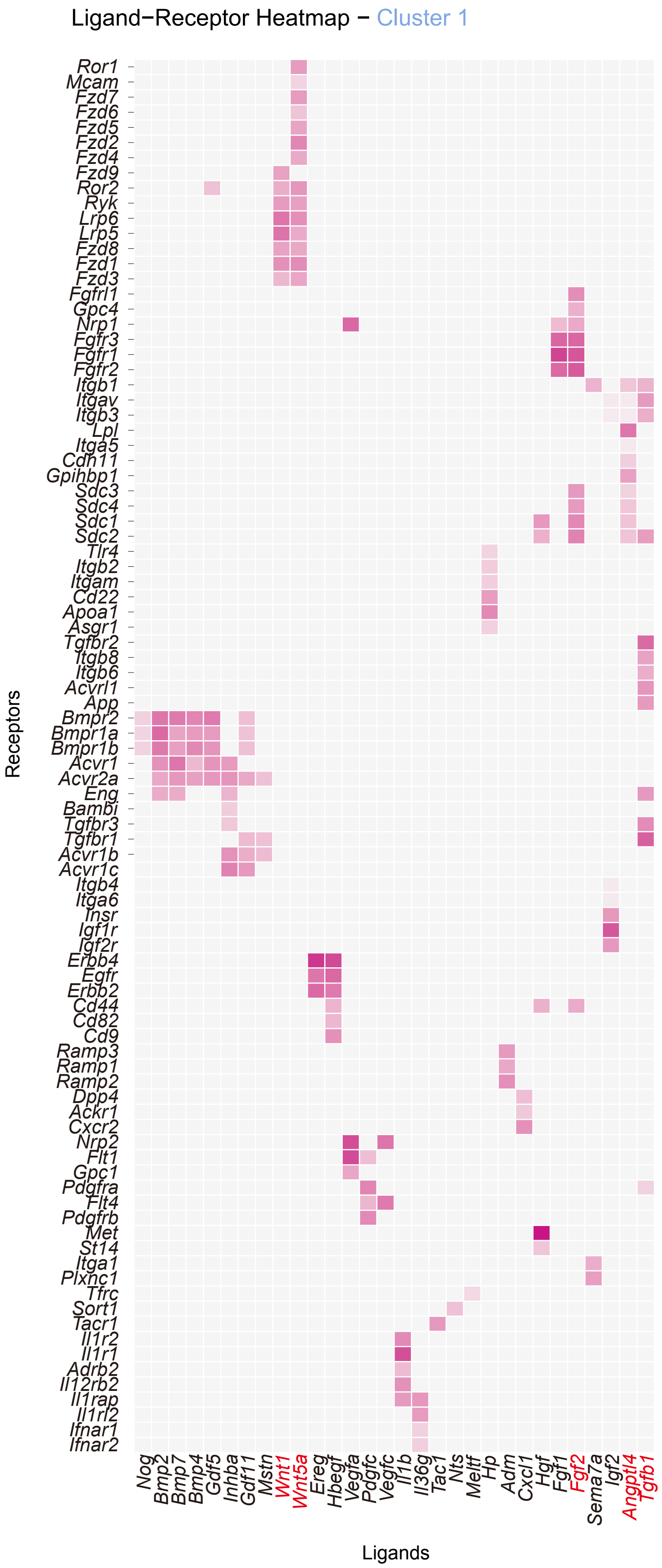


**Figure S11 Predicted signaling in the matrix-enriched subpopulation (Seurat cluster 1) in *Mepe*$^{Cre/+}$-sorted osteocytes.**
Heatmap of predicted ligands (signal) and receptors activated in the terminative matrix-enriched cluster (cluster 1) are shown. Ligands highlighted in red correspond to the top five receptors that express more numbers of receptors in the clusters.

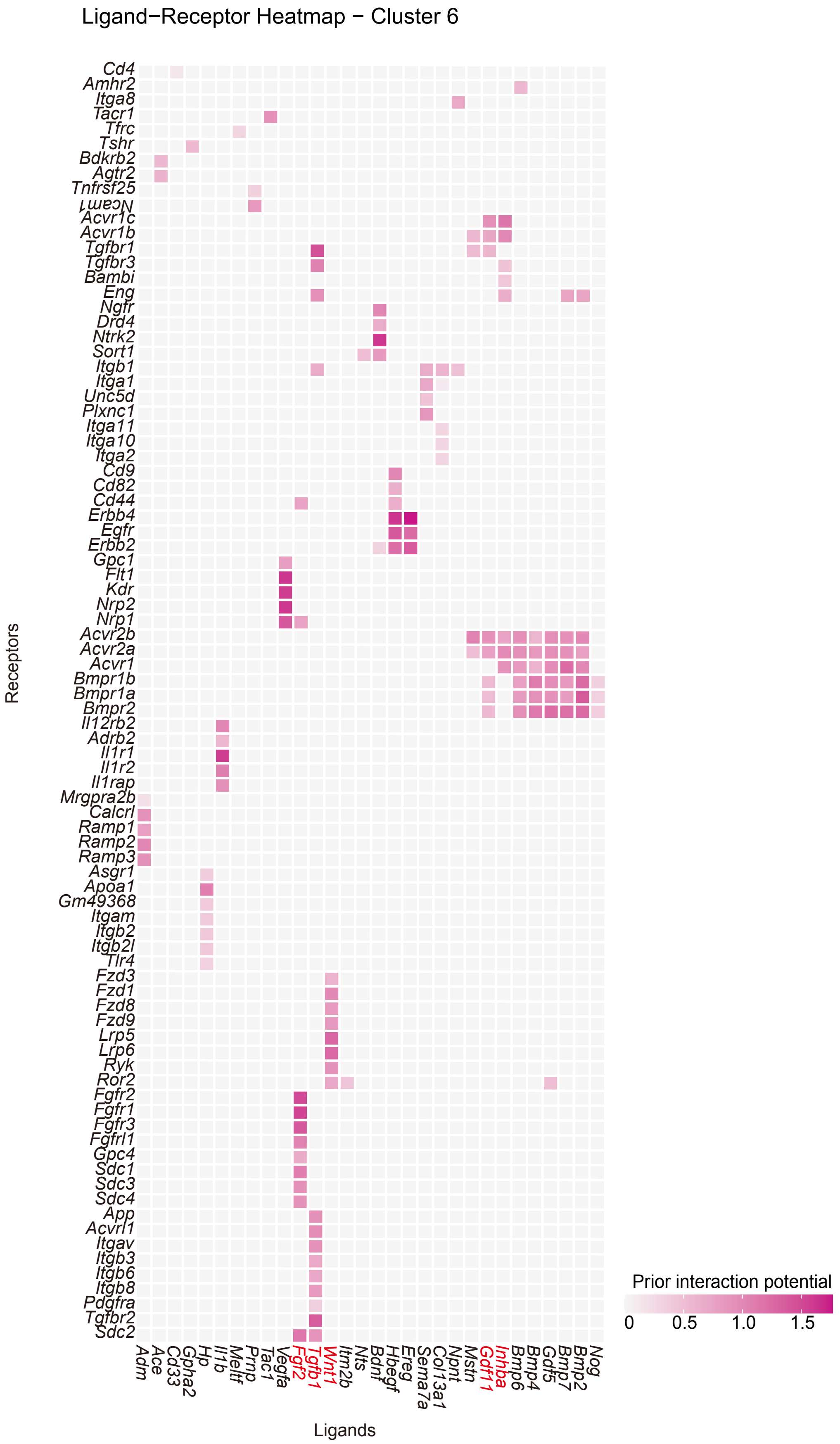


**Figure S11 Predicted signaling in the matrix-enriched subpopulation (Seurat cluster 6) in *Mepe*[Cre/+]-sorted osteocytes.**
Heatmap of predicted ligands (signal) and receptors activated in the terminative matrix-enriched cluster (cluster 6) are shown. Ligands highlighted in red correspond to the top five receptors that express more numbers of receptors in the clusters.

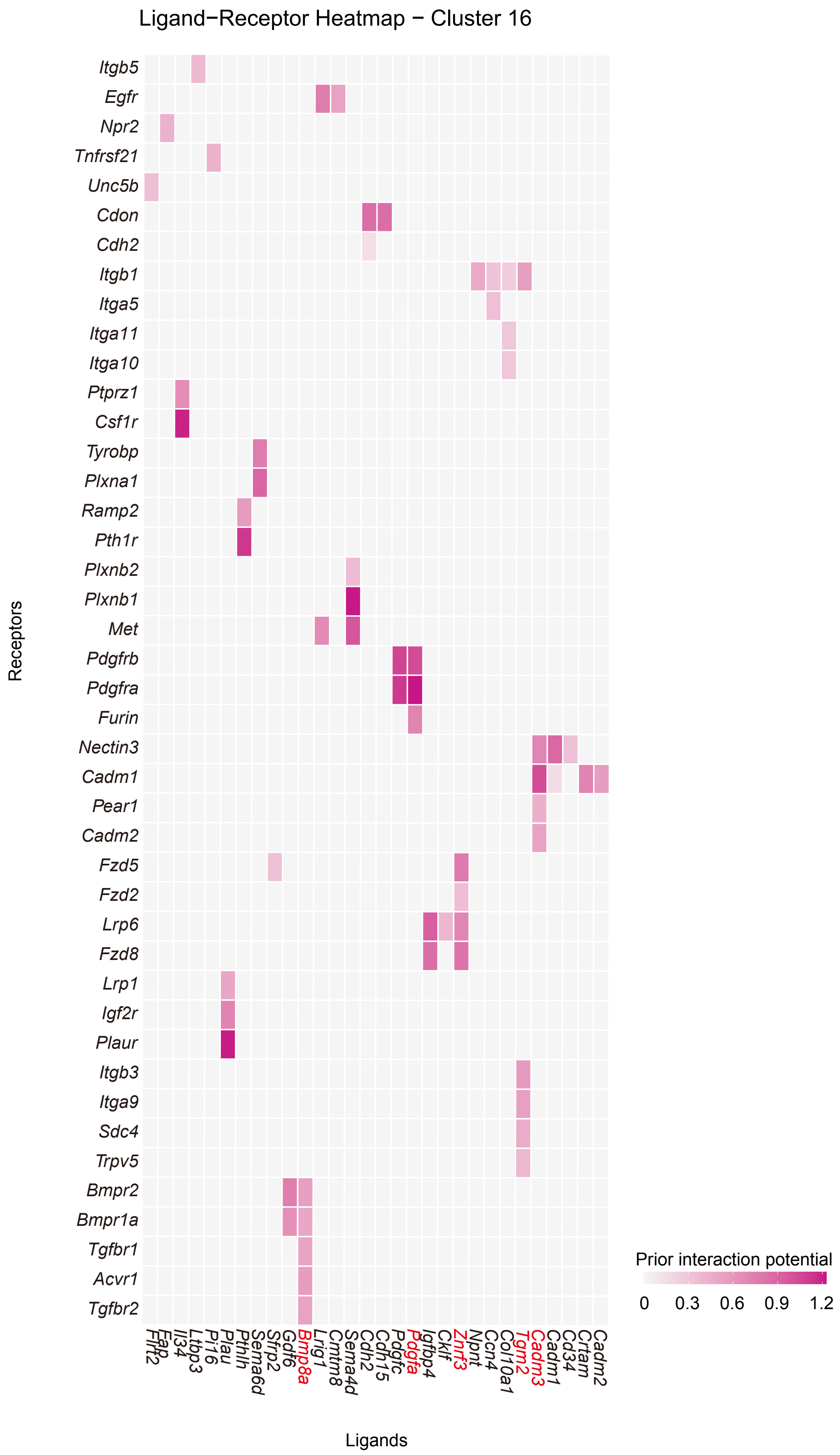


**Figure S12 Predicted signaling in the matrix-enriched subpopulation (Seurat cluster 16) in *Mepe*$^{Cre/+}$-sorted osteocytes.**
Heatmap of predicted ligands (signal) and receptors activated in the terminative matrix-enriched cluster (cluster 16) are shown. Ligands highlighted in red correspond to the top five receptors that express more numbers of receptors in the clusters.

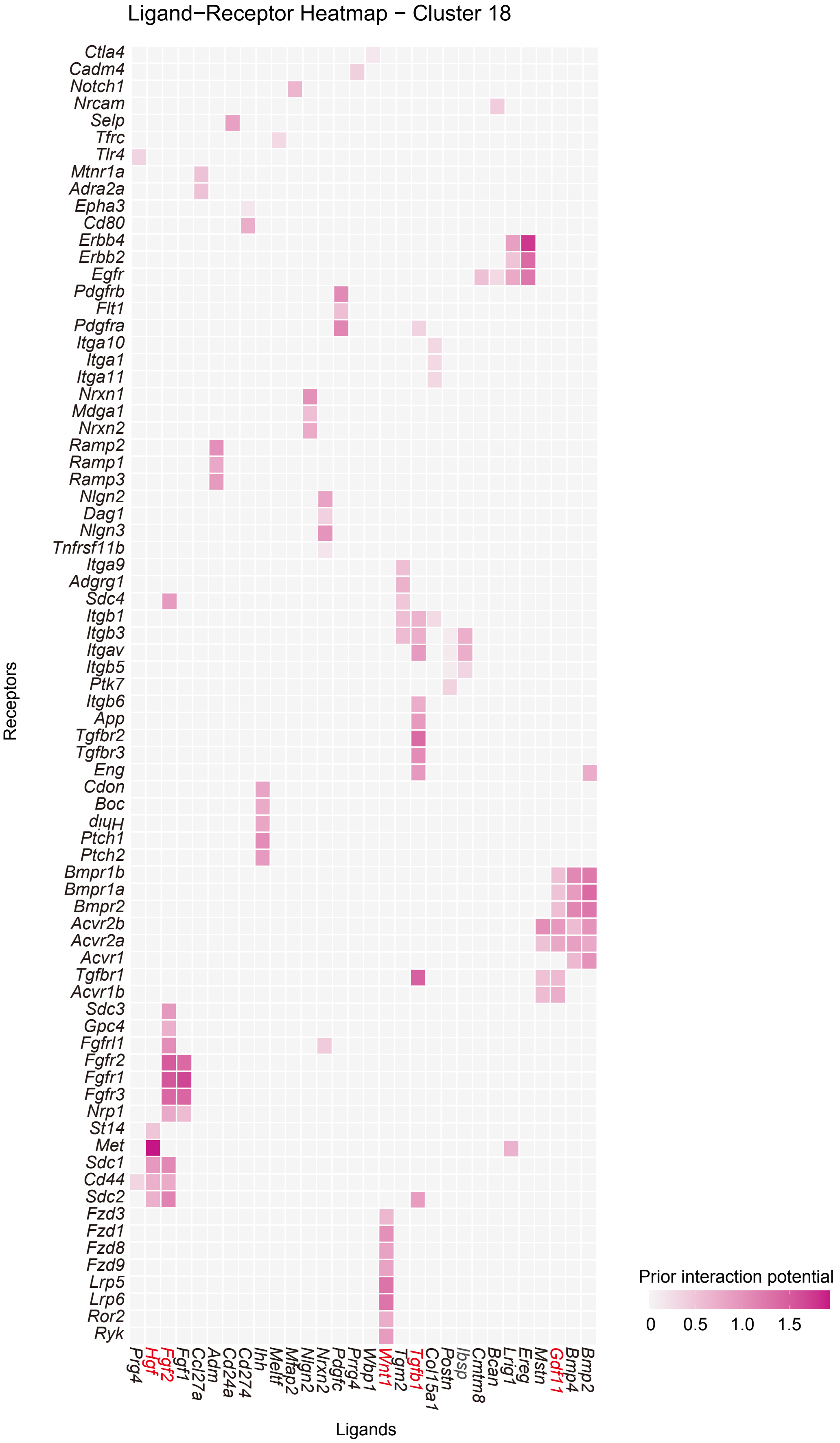


**Figure S13 Predicted signaling in the matrix-enriched subpopulation (Seurat cluster 18) in *Mepe*Cre/+-sorted osteocytes.**
Heatmap of predicted ligands (signal) and receptors activated in the terminative matrix-enriched cluster (cluster 18) are shown. Ligands highlighted in red correspond to the top five receptors that express more numbers of receptors in the clusters.

**A**

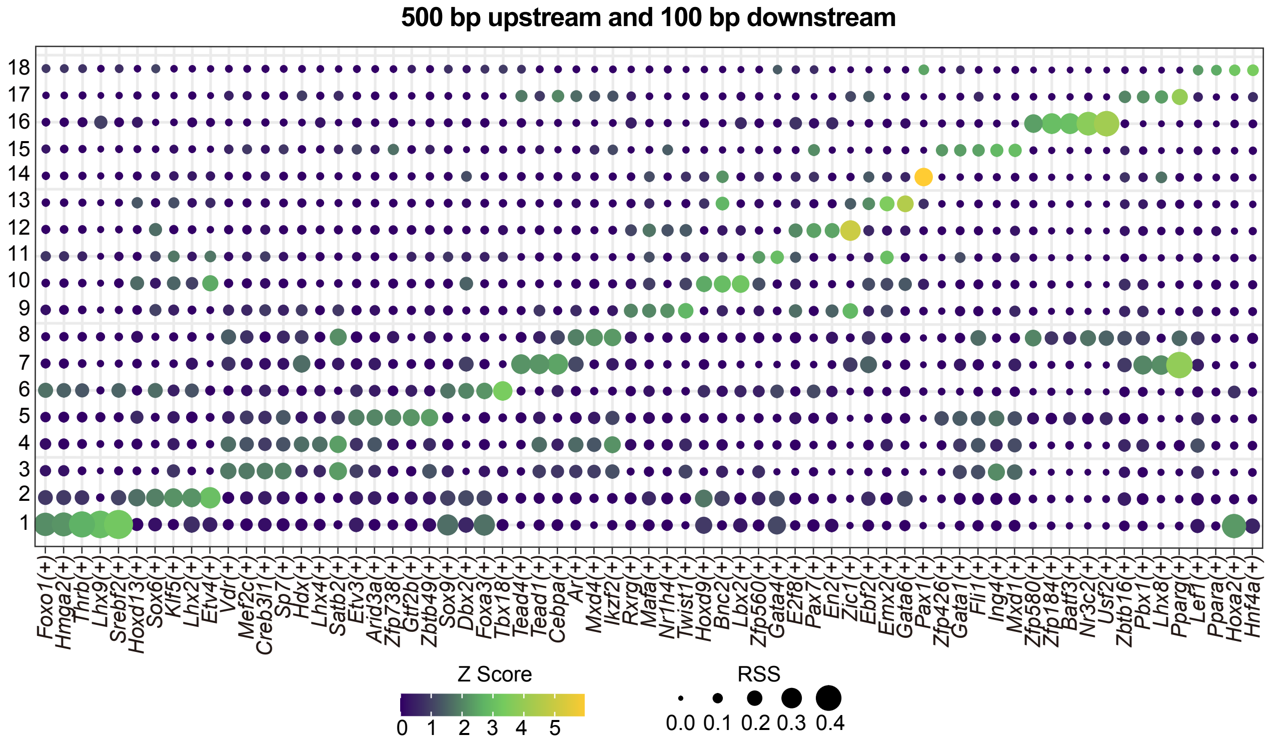


**B**

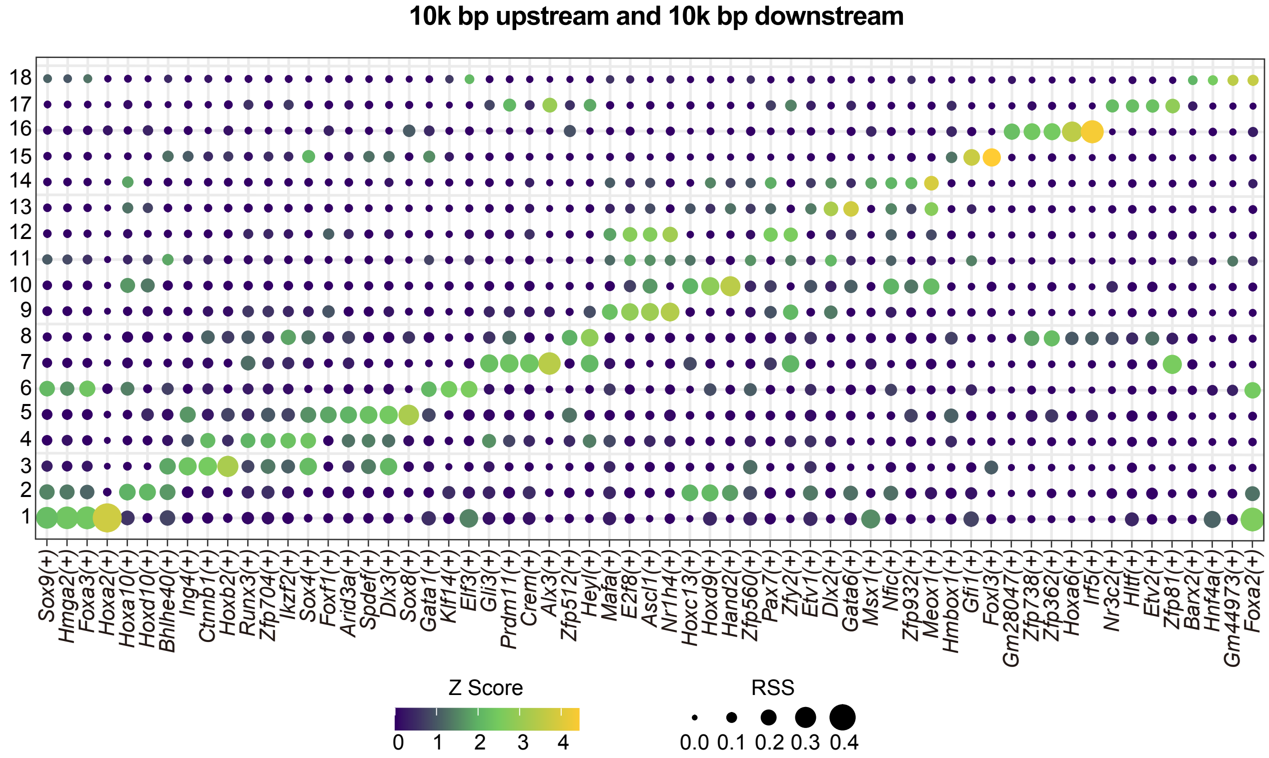


**Figure S14 Evaluation of potential transcription factors in the maturing process of *Mepe*$^{Cre/+}$-sorted osteocytes.**
**(A, B)** Possible transcription factors in a narrow and broad range were evaluated by pySCENIC.

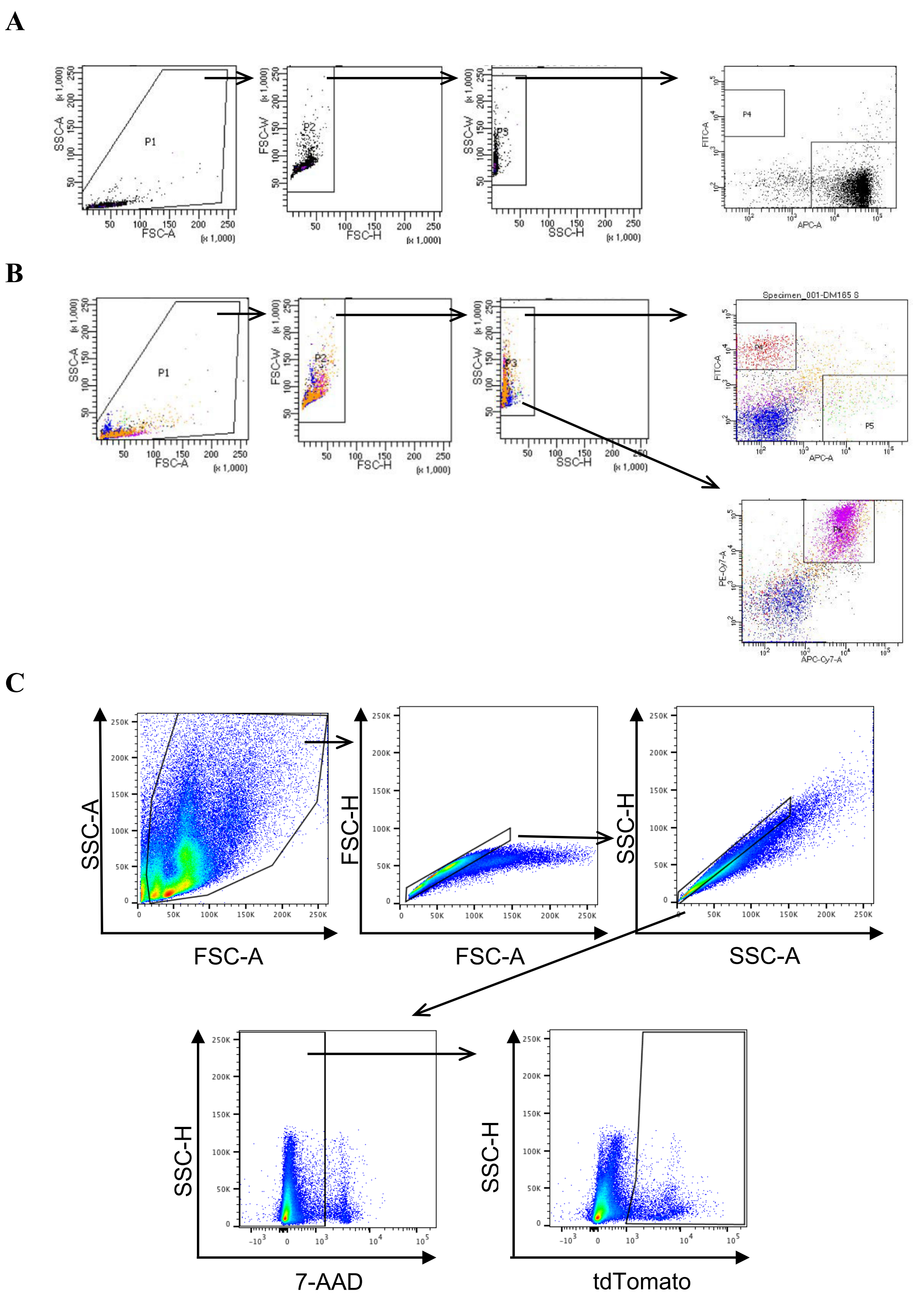


**Figure S15 Gating strategies of flowcytometric analyses.**

**(A)** Thymus immune cells analysis strategy. FITC channel represents for CD4 and APC channel represents for CD8a. (**B)** Spleen immune cells analysis strategy. FITC channel represents for CD3, APC channel represents for CD11b, PE-Cy7 channel represents for B220, and APC-Cy7 channel represents for CD19. (**C)** Osteocytes sorting strategy.

**Table S1 Serum biochemical profiles of osteocyte-ablated mice.**

Young mice (10-weeks-old)

| | Control | Ablation | P-value | t-test |
|---|---|---|---|---|
| Ca (mEq/L) | 11.39 ± 0.04041 | 11.2 ± 0.1704 | 0.3099 | ns |
| P (mEq/L) | 9.143 ± 0.2379 | 8.657 ± 0.5291 | 0.4188 | ns |
| TP (mEq/L) | 6.043 ± 0.1232 | 6.314 ± 0.1010 | 0.1141 | ns |
| ALB (mEq/L) | 3.829 ± 0.07143 | 3.843 ± 0.06117 | 0.7161 | ns |
| AST (mEq/L) | 137.3 ± 22.92 | 176.3 ± 40.40 | 0.1934 | ns |
| ALT (mEq/L) | 37.57 ± 7.023 | 33.43 ± 3.108 | 0.5995 | ns |
| BUN (mEq/L) | 31.53 ± 1.743 | 29.04 ± 1.306 | 0.276 | ns |
| CRE (mEq/L) | 0.1071 ± 0.008081 | 0.1000 ± 0.004880 | 0.4639 | ns |

Aged mice (52-weeks-old)

| | Control | Ablation | P-value | t-test |
|---|---|---|---|---|
| Ca (mEq/L) | 11.54 ± 0.2015 | 11.30 ± 0.1225 | 0.3386 | ns |
| P (mEq/L) | 10.84 ± 0.6969 | 10.14 ± 0.4718 | 0.4296 | ns |
| TP (mEq/L) | 6.720 ± 0.2354 | 6.360 ± 0.1749 | 0.2545 | ns |
| ALB (mEq/L) | 4.040 ± 0.1208 | 3.780 ± 0.09165 | 0.1248 | ns |
| AST (mEq/L) | 125.4 ± 18.97 | 114.6 ± 15.17 | 0.6684 | ns |
| ALT (mEq/L) | 42.60 ± 6.705 | 33.60 ± 6.592 | 0.3665 | ns |
| BUN (mEq/L) | 25.74 ± 1.407 | 28.20 ± 0.5621 | 0.1431 | ns |
| CRE (mEq/L) | 0.1000 ± 0.006325 | 0.09800 ± 0.005831 | 0.822 | ns |

Seven (young) and five (aged) biological replicates were used for the measurement of serum biochemical profiles. mEq, mili equivalent; ns, statistically non-specific. Statistical analysis was performed by two-tailed unpaired t-test with Welch's correction.

**Text S1 Generation of mice carrying the *Mepe-IRES-Cre* allele**.

All fragments for the construction of the *Mepe-IRES-Cre* targeting vector were derived from a BAC clone (RP23-150D15: BACPAC Resource Center) by restriction digestion and subcloned into a pBluescript II vector. The homology arms were cloned into a targeting vector backbone containing *PGK-neo* and *IRES-Cre* alle (IRES-Cre-pA FRT PGK neo bpA FRT vector; Fig. S1A). The long arm was excised as a EcoRI/SpeI fragment (5.3 kb) from the pBluescript II vector including the first half homology arms that contains CDS part of exon 3 of *Mepe*, and cloned into the IRES-Cre-pA FRT PKG neo bpA FRT vector upstream of the *IRES-Cre* alle. The short arm that contains 3'-UTR part of exon 3 of *Mepe* was excised as the SpeI/ PciI fragment (0.8 kb), then cloned into the downstream of *FRT* flanked *PGK-neo* alle. The targeting vector was linearized and electroporated into mouse embryonic stem (ES) cells. After 8–9 days of G418 selection, resistant single colonies were picked and transferred onto primary embryonic fibroblast cells acting as feeder cells, and expanded to allow diagnosis of homologous recombination. Targeted ES cells were injected into C57BL/6J blastocysts to achieve initial germ-line transmission. Chimeric mice were crossed with C57BL/6J to establish a line for the *Mepe-IRES-Cre-Neo* allele. Mice carrying the *FRT*-flanked *PGK-neo* cassette was crossed with C57BL/6J background *FLPe* transgenic mice to excise the *PGK-neo* cassette and subsequently crossed to C57BL/6J mice to remove the *FLPe* recombinase transgene to generate *Mepe-IRES-Cre* mice.

Potential recombinant mice were detected by genomic long PCR (Fig. S1B). The forward primer (P1, AGTGGGAAGGATGTTTGCAT) binds within the intron region before exon 3, the reverse primer (P2, CTCCCCGTCCTCTAACTGAAG) binds to a unique region outside the short arm of the targeting vector and within the endogenous locus (Fig. S1B). Routine gene typing was performed by PCR using the primers (P3, CTCGTGCTTTACGGTATC; P4, TGTTGGCTTGCTCAGTTC).
